# Epigenome erosion drives neural crest-like phenotypic mimicry in triple-negative breast cancer and other SOX10+ malignancies

**DOI:** 10.1101/2021.03.30.437624

**Authors:** Jodi M. Saunus, Xavier M. De Luca, Korinne Northwood, Ashwini Raghavendra, Alexander Hasson, Amy E. McCart Reed, Malcolm Lim, Samir Lal, Ana Cristina Vargas, Jamie R. Kutasovic, Andrew J. Dalley, Mariska Miranda, Emarene Kalaw, Priyakshi Kalita-de Croft, Irma Gresshoff, Fares Al-Ejeh, Julia M.W. Gee, Chris Ormandy, Kum Kum Khanna, Jonathan Beesley, Georgia Chenevix-Trench, Andrew R. Green, Emad A. Rakha, Ian O. Ellis, Dan V. Nicolau, Peter T. Simpson, Sunil R. Lakhani

## Abstract

**Background:** Intratumoural heterogeneity is a poor prognostic feature in triple-negative breast cancer (TNBC) and other high-grade malignancies. It is caused by genomic instability and phenotypic plasticity, but how these features co-evolve during tumour development remains unclear. SOX10 is a transcription factor, neural crest stem cell (NCSC) specifier and candidate mediator of cancer-associated phenotypic plasticity.

**Methods:** Using immunophenotyping, we investigated the expression of SOX10 in normal human breast tissue and breast cancer (n=21 cosmetic breast reduction and 1,860 tumour samples with clinical annotation). We then defined the context and evolution of its expression in TNBC compared to 21 other malignancies using systems-level transcriptomics.

**Results:** SOX10 was detected in nuclei of normal mammary luminal progenitor cells, the histogenic origin of most TNBCs. In breast cancer, nuclear SOX10 predicted poor outcome amongst cross-sectional (log-rank p=0.0015, hazard ratio 2.02, n=224) and metaplastic (log-rank p=0.04, n=66) TNBCs. Systems-level transcriptional network analysis identified a core module in SOX10’s normal mammary epithelial transcription program that is rewired to NCSC genes in TNBC. Reprogramming was proportional to DNA damage and genome-wide promoter hypomethylation, particularly at CpG island shores. Using a novel network analysis pipeline, we found that NCSC-like transcriptional reprogramming is also strongly associated with promoter hypomethylation in other SOX10+ malignancies: glioma and melanoma.

**Conclusions:** We propose that cancer-associated genome hypomethylation simulates the open chromatin landscape of more primitive cell states, and that on this relatively unrestricted background, SOX10 recreates its ancestral gene regulatory circuits by default. These findings provide new insights about the basis of intratumoural heterogeneity and resurrection of developmental phenotypes in cancer; and highlight the potential for therapeutics that limit chromatin remodelling.

## Introduction

Effective management of triple-negative breast cancer (TNBC) remains a significant challenge worldwide. These tumours lack expression of oestrogen and progesterone receptors (ER/PR) and HER2, hence are not indicated for treatment with classical molecular-targeted agents. Chemotherapy remains the most reliable systemic treatment option, producing durable responses in ~60% of patients, while the other ~40% typically present with lung, liver and/or brain metastases within five years^1–3^. Second-line chemotherapy can temporarily stabilise metastatic disease but is rarely curative, so these patients endure a heavy treatment burden for no lasting benefit. Efforts to develop alternative treatments have been hampered by molecular and cellular variability between, and within, individual tumours. Intra-tumoural heterogeneity (ITH) directly increases the probability of relapse because it diversifies the substrate for clonal selection^4–7^. It has been proposed that to further improve the prognosis for TNBC patients, we need to develop agents that target the drivers of heterogeneity itself^8^.

TNBCs are characterised by defective DNA repair, mitotic spindle dysfunction, chromosomal aberrations and a mutation rate approximately 13 times that of other breast tumours^4,5^. Genomic instability is a key driver of ITH, however only some cases can be explained by selection of individual driver mutations^9^, and other sources of heterogeneity are coming to light^10–12^. For example, cellular heterogeneity is influenced by the differentiation state of the normal cellular precursor(s)^13^, which in TNBC is thought to be luminally-committed progenitor (LP) cells^14–17^.

ITH is also driven by phenotypic plasticity – the dynamic reprogramming of cell state in response to extrinsic stimuli^10,11^. Cancer cell state transitions can be de-differentiating (the loss of lineage commitment and acquisition of stem cell features) and/or trans-differentiating (assuming the state of another cell type)^18^. Compared to genomic and histogenic sources of ITH, how tumour cells invoke this capability is poorly understood, and yet potentially more ominous for the patient, as cell state transitions can be induced by treatment via heritable epigenetic change. In controlled experimental conditions, drug tolerant TNBC cell states can be averted by epigenome remodelling inhibitors^19–23^, suggesting these agents might reduce rates of relapse if used clinically^8,11^. However, epigenetic therapies have genome-wide effects, so our ability to use them rationally requires a deeper understanding of the epigenome-driven features that characterise treatment-refractory human tumours^8^.

SOX10 is a transcription factor that was recently implicated in mediating phenotypic plasticity in experimental models of TNBC^24^. It is first expressed in embryonic neural crest stem cells (NCSCs), where its self-reinforcing gene regulatory module facilitates multipotency and cell migration, orchestrating the embryo patterning process^25–28^. Once patterning is complete, *SOX10* is silenced in all NCSC descendants except glial and melanocyte progenitors; and is nascently induced in ectoderm-derived epithelial progenitor cells of the salivary, lacrimal and mammary glands^29–33^. In contrast to NCSCs where the genome is widely unmethylated and accessible for transcription, SOX10’s regulation of self-renewal and differentiation in adult tissues is determined by the lineage-specific epigenetic fingerprint. Remarkably, ectopic expression of *SOX10* reprogrammed postnatal fibroblasts with multipotency and migration capabilities equivalent to NCSCs, providing they were also exposed to chromatin unpacking agents and early morphogens (DNA methylation and histone deacetylase inhibitors plus Wnt and BMP agonists)^34^. This established that with the erasure of lineagespecific epigenetic marks and appropriate extrinsic cues, SOX10 recreated its ‘default’ regulatory circuit, and that this is sufficient to phenocopy NCSCs.

SOX10 expression in human breast cancer is associated with TN, basal-like, metaplastic and neural progenitor-like phenotypes^4,35–39^. In transgenic mouse mammary tumour cells, it promoted invasiveness, expression of mammary stem/progenitor, EMT and NCSC genes and the repression of epithelial differentiation genes^24^. These findings suggest that SOX10 could mediate de-differentiation in TNBC; but the clinical relevance is unclear, particularly given there are no available inhibitors of SOX10 itself. We explored the significance of SOX10 in breast cancer development and progression by immunophenotyping histologically normal breast tissue, and large breast tumour sample cohorts. To understand its contribution to phenotypic plasticity and identify drivers of this capability, we performed systems-level analysis of breast cancer omics datasets to map SOX10’s regulatory circuit in the broader TNBC transcriptional network.

## Materials, methods and data availability

See supplementary file 1.

## Results

### SOX10 is expressed in luminal progenitor cells of the human mammary gland

To confirm the expression pattern of SOX10 in the human breast, we performed immunohistochemical (IHC) analysis of 19 histologically normal reduction mammoplasty (RM) samples using a validated antibody (Fig-S1a, Table-S1). SOX10 was detected in the nuclei of ductal and lobular epithelia, with individual terminal ducto-lobular units (TDLUs) exhibiting either basal-restricted or combined baso-luminal expression (Fig-1a). Compared to ducts, lobules were more likely to exhibit luminal compartment expression of SOX10 (Fig-1b), consistent with a role in lobulogenesis. Indeed, TDLUs with basal-restricted SOX10 expressed high levels of luminal cytokeratins (CK)8/18, while TDLUs with dual-compartment SOX10 had low CK8/18. This was evident even in neighbouring structures of the same specimen (Fig-1c, Fig-S1b). IHC analysis of serial sections showed SOX10+ luminal cells lacked ER and were positive for the LP marker c-kit, with no obvious relationship to proliferation marker Ki-67 (Fig-1d). We also analysed *SOX10* mRNA in a published dataset from FACS-sorted human mammary epithelial cells (hMECs)^15^. *SOX10* levels were similar to established LP markers *ELF5* and *KIT:* highest in EpCAM+/CD49f+ LP cells, moderate in the EpCAM-/CD49f+ basal compartment (myoepithelia and mammary stem cells (MaSCs)) and low in EpCAM+/CD49f-mature luminal (ML) cells (Fig-1e).

**Figure-1.**
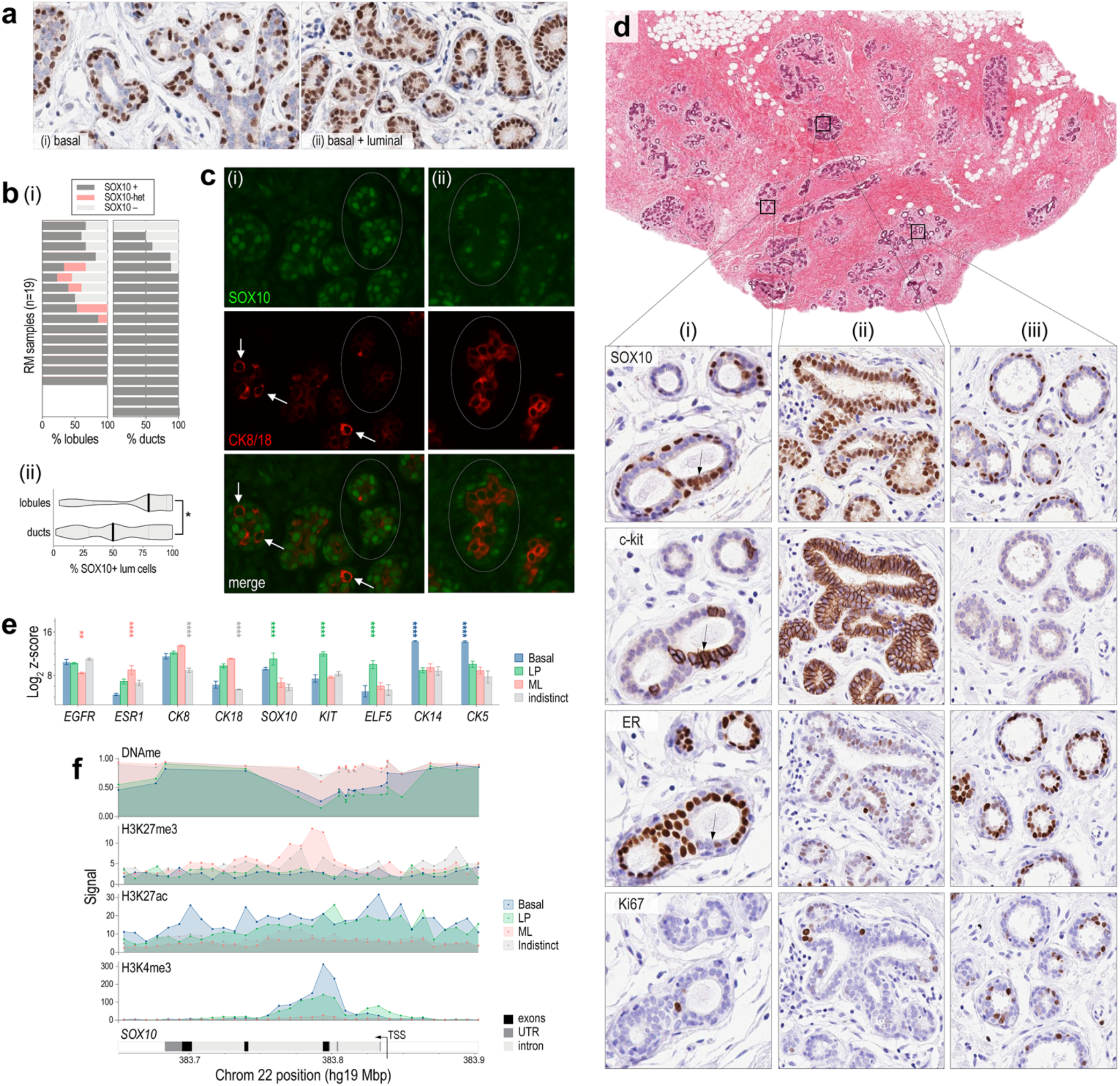
SOX10 is expressed in basal and luminal progenitor cells of the human mammary gland. (**a**) Representative SOX10 IHC analysis of reduction mammoplasty (RM) samples. Some terminal ducto-lobular units (TDLUs) had exclusive basal compartment expression (i) while others had expression in both basal and luminal compartments (ii). (**b**)(i) Analysis of SOX10 expression in ducts vs lobules of RM samples from 19 donors (whole sections). (ii) SOX10 expression in lobules was heterogeneous and more likely to occur in the luminal compartment (Mann-Whitney p=0.011; n=102 ducts and 102 lobules; median +/− 95% confidence interval shown). (**c**) Representative immunofluorescent staining of SOX10 and CK8/18. Circled lobules and isolated cells (arrows) exhibited reciprocal expression of SOX10 (green) and CK8/18 (red) in structures with either (i) dual compartment (ii) or basal-restricted SOX10 expression. (**d**) IHC analysis of SOX10, c-kit, ER and Ki67 in serial RM sections. The three magnified regions represent major SOX10 staining patterns: (i) dual compartment, heterogeneous; (ii) dual compartment, homogeneous; and (iii) basal-restricted. Luminal SOX10 expression was directly associated with c-kit and inversely associated with ER, with no obvious relationship to Ki67 (e.g., cell cluster indicated with an arrow). (**e**) *SOX10* mIRNA levels in FACS-sorted human mammary epithelial cell (hMEC) subtypes^15^. Differentiation markers were analysed for comparison: basal markers CK14 and CK5; luminal progenitor (LP) markers *KIT* and *ELF5;* and markers enriched in mature luminal (ML) cells: *CK18* and *ESR1* (isolates with significantly different marker levels according to paired ANOVA tests are indicated and colour-coded: \*\*\*\**p*<0.00001; \*\*\**p*<0.0001; \*\**p*<0.001). (**f**) Average Illumina EPIC 850k methylation array beta values of *SOX10* probes in FACS-sorted hMEC samples (DNAme), aligned with histone modification signals in a published ChIP-seq dataset^42^: H3K4me3, H3K27ac (activating) and H3K27me3 (repressive). Data are represented to scale on human chromosome 22. *TSS, transcription start site; UTR, untranslated region.* Indistinct = negative for CD45 (hematopoietic cells), CD31 (endothelia), CD235a (erythrocyte precursors), EpCAM and CD49f (epithelia).

SOX10 is epigenetically regulated in mouse mammary gland^40,41^, so we investigated this in human tissue. We isolated hMECs from two fresh RM samples using FACS with antibodies against CD49f and EpCAM, then performed high-density DNA methylation array profiling. *SOX10* was hypomethylated in LP and basal samples (*p*<1.0E^-06^; Fig-1f). Consistently, analysis of hMEC chromatin immuno-precipitation sequencing (ChIP-seq) data from six independent RM samples^42^ showed the *SOX10* locus is enriched with activating (H3K4me3, H3K27ac) and depleted of repressive H3K27me3 marks in LP and basal samples (Fig-1f).

### SOX10 is associated with poor clinical outcomes in TNBC

Analysis of TCGA, METABRIC and ICGC breast tumour datasets^43–45^ showed *SOX10* mRNA is expressed almost exclusively in TNBC, with a bimodal distribution suggesting distinct SOX10 positive and negative (+/-) subgroups (Fig-2a, Fig-S2a). Copy-number (CN) amplification or gain at the *SOX10* locus was evident in ~20% of TNBCs (Fig-2b) and was associated with higher mRNA levels in both METABRIC and TCGA datasets (Fisher’s Exact *p*≤0.001). Analysis of TCGA HM450k methylation array data indicated that *SOX10* is frequently hypomethylated in TNBC (Fig-2b), and that this correlates strongly with expression (Fig-2c, Figs S2c-d), but does not extend to adjacent genes on chromosome 22 (Fig-2d). Hence, positively selected CN gains and gene-specific hypomethylation underpin *SOX10* expression in a subset of TNBCs.

**Figure-2.**
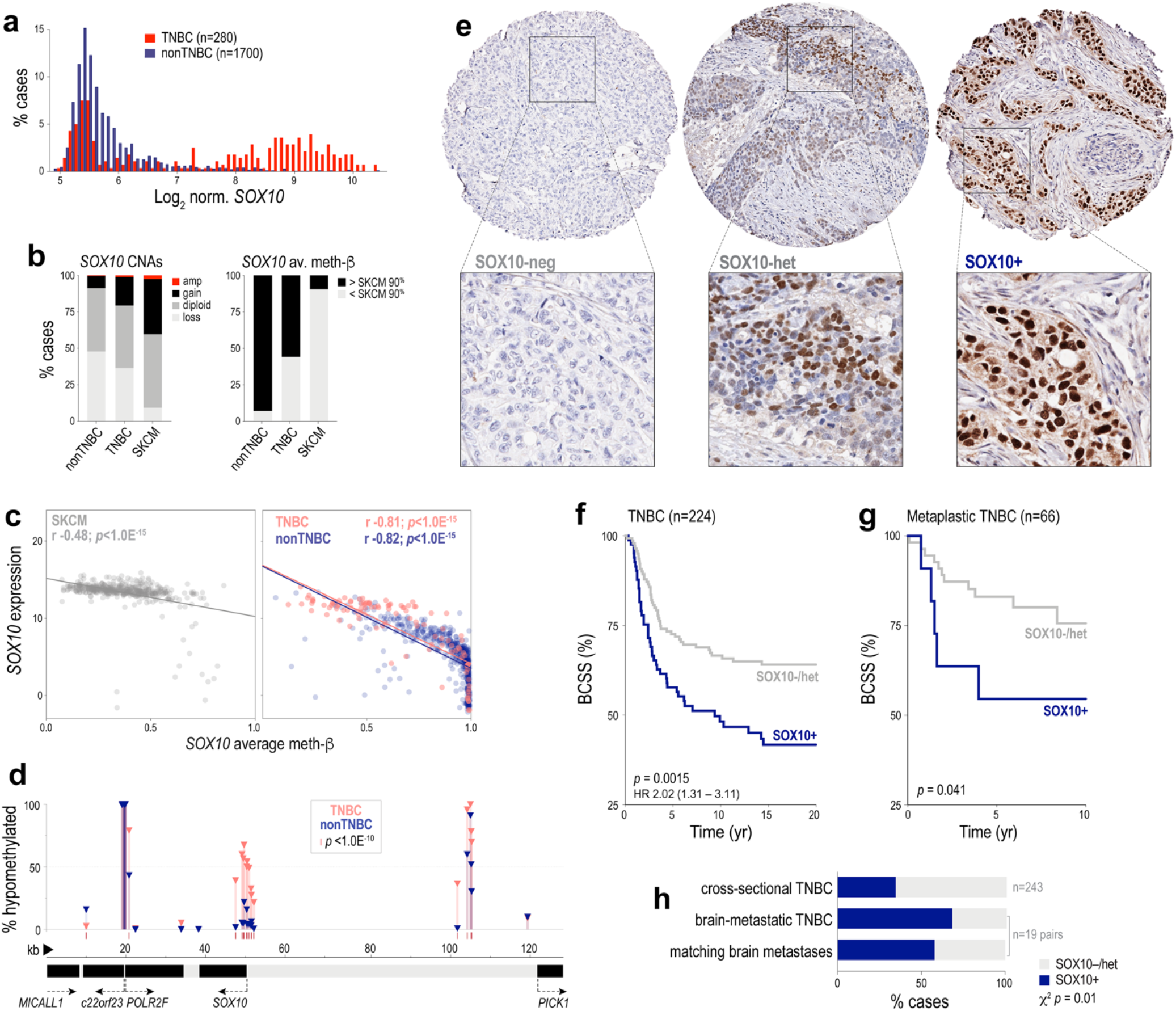
Expression of SOX10 in human breast cancer. (**a**) Bimodal expression of *SOX10* in TNBC compared to other breast cancers (nonTNBC) in the METABRIC cohort. (**b**) Frequency of copy-number alterations (CNAs) and DNA hypomethylation affecting *SOX10* in TNBC and nonTNBC compared to the archetypal SOX10+ malignancy, melanoma (SKCM) (TCGA datasets). (**c**) Correlation between *SOX10* methylation and expression (normalised RNAseq counts) in SKCM, TNBC and nonTNBC (Spearman correlation coefficients (r) and *p*-values are shown; derived from TCGA data). (**d**) Proportions of TNBC and nonTNBC cases with hypomethylation at each probe across the *SOX10* locus (as defined in (b)). (**e**) Representative IHC showing SOX10-neg, heterogeneous and nuclear-positive (+) TNBCs. Tumours with absent or very weak nuclear staining in ≥50% of tumour cells were classified as SOX10-negative, while those with any one of replicate TMA cores exhibiting moderate-strong nuclear staining in <50% OR weak-moderate nuclear staining in ≥50% of tumour cells were classified as heterogeneous (see also Fig-S2g). Survival curves of heterogeneous and negative categories overlapped (Fig-S2i) hence are grouped together here. (**f**) Kaplan Meier analysis of the relationship between SOX10 nuclear positivity and breast cancer-specific survival (BCSS) in cross-sectional TNBCs. Log-rank test *p-*value and hazard ratio (HR) shown (95% confidence interval)). (**g**) Kaplan Meier analysis of the relationship between SOX10 nuclear positivity and BCSS in TNBCs classified as metaplastic breast cancers. Gehan-Breslow-Wilcoxon test *p-*value shown. (**h**) SOX10 expression in brain metastatic TNBC and matching brain metastases (BrM), compared to the frequency in cross-sectional TNBCs (Chi-square *p*-value shown).

Analysing published cell line gene expression and methylation array datasets^46,47^ and our cell line bank^48,49^, we found that in contrast to tumours, TNBC cell lines express very low to undetectable levels of *SOX10,* and the *SOX10* gene is hypermethylated (Figs S2d-e). shRNA-mediated depletion of *SOX10* in one of the few positive lines (HCC1569) resulted in 100% cell death within a few passages (Fig-S2f).

Next, we IHC studies to investigate the prognostic significance of SOX10 expression at the protein level. Surveying a large, cross-sectional cohort of invasive breast tumours from Australia and the UK (n=1,330), we detected SOX10 almost exclusively in tumour cell nuclei of TN cases (Fig-2e). Approximately 38% of TNBCs were classified as SOX10+, and another 11.5% exhibited heterogeneous staining (Fig-2e, Fig-S2g). SOX10 positivity was associated with histologic features typical of this group, such as high grade, metaplastic and medullary morphology, pushing margins and larger size at diagnosis (Table-S2). Similar, though statistically weaker trends were found between these variables and heterogeneous SOX10 staining (Fig-S2h).

Rather than a simple correlate of the TN phenotype, SOX10 positivity stratified TNBC-specific survival in both univariate (Fig-2f, Fig-S2i) and multivariate regression analyses, with prognostic value greater than clinicopathologic indicators used in current clinical practice: tumour size, grade and the density of tumour-infiltrating lymphocytes (TILs) (hazard ratio 1.8-2.5; *p*=0.02-0.002; Table-S2). Increased propensity for brain metastasis is one of the factors underlying premature death in TNBC, so we performed IHC analysis of retrospectively identified brain metastatic TNBCs and their matching brain metastases (n=19 pairs). Compared to cross-sectional TNBCs, SOX10 was over-represented in the brain-metastatic cases, and SOX10 status was concordant in ~90% of matching brain tumours (Fig-2h).

Consistent with previous reports^37,50^, we also detected nuclear SOX10 in an independent cohort of metaplastic breast cancers (MBC; from the Asia-Pacific Metaplastic Breast Cancer consortium^51^). Compared to cross-sectional cases, SOX10 staining was more heterogeneous in the MBC cohort and was not associated with TN status (Fig-S2j); but was prognostic in MBCs with a TN phenotype (Fig-2g).

Considering all of our IHC study findings, we concluded that strong nuclear expression of SOX10 is associated with TNBC progression.

### SOX10’s TNBC regulatory module confers transcriptomic similarity to NCSCs

To investigate the basis of SOX10’s association with poor patient outcomes, we compared the expression profiles of TNBCs expressing high versus low levels of *SOX10* mRNA and found that *SOX10^high^* tumours were significantly enriched with the expression of mesenchymal, neural and glial development genes (Fig-S3, Tables S3-S4).

We then mapped *SOX10’s* regulatory neighbourhood within the breast cancer transcriptome using weighted gene co-expression network analysis (WGCNA). This approach quantifies co-variation in gene expression across a biological sample set to identify genes with highly coordinated regulation, which is indicative of functional relatedness^52,53^. We built a network from TCGA breast cancer RNAseq data (n=919 cases) and validated it with datasets from METABRIC (n=1278, expression array) and ICGC (n=342, RNAseq). In this model, all genes expressed above a background threshold are connected (12,588 genes, 12,588^2^ connections). The connection between each gene pair is based on a weighted correlation coefficient, and unsupervised clustering can reveal groups of genes with a high probability of co-functionality (modular transcription programs). The module eigengene (ME) is a centroid calculated for each module in each sample that represents both module expression and net connection strength.

WGCNA partitioned ~20% of expressed genes into eight consensus modules that align with established hallmarks of breast cancer; for example, an ER/FOXA1-driven module expressed in luminal tumours, and a mitotic instability module in basal-like and luminal-B tumours (Table-1, Fig-3a, Tables S5-S8, Supp file-2). The remaining ~80% of genes were not linked to any one module in particular. *SOX10* was identified as one of the most interconnected genes in the ‘green’ module, which has a hierarchical structure (Figs S4a-b) and is predominantly expressed in high-grade TNBCs (Fig-S4c). In this module, SOX10’s co-expression profile was highly similar to genes implicated in Wnt signalling, neuroglial differentiation and embryo patterning (Fig-3b). We named it the SOXE-module and ascribed ‘multipotency’ as its primary ontology, as the member gene list is enriched with developmental phenotypes, includes all three SOXE family members *(SOX8/9/10)* and embryonic stem cell genes *(LMO4, POU5F1)* (Fig-3c, Table-S9).

**Figure-3.**
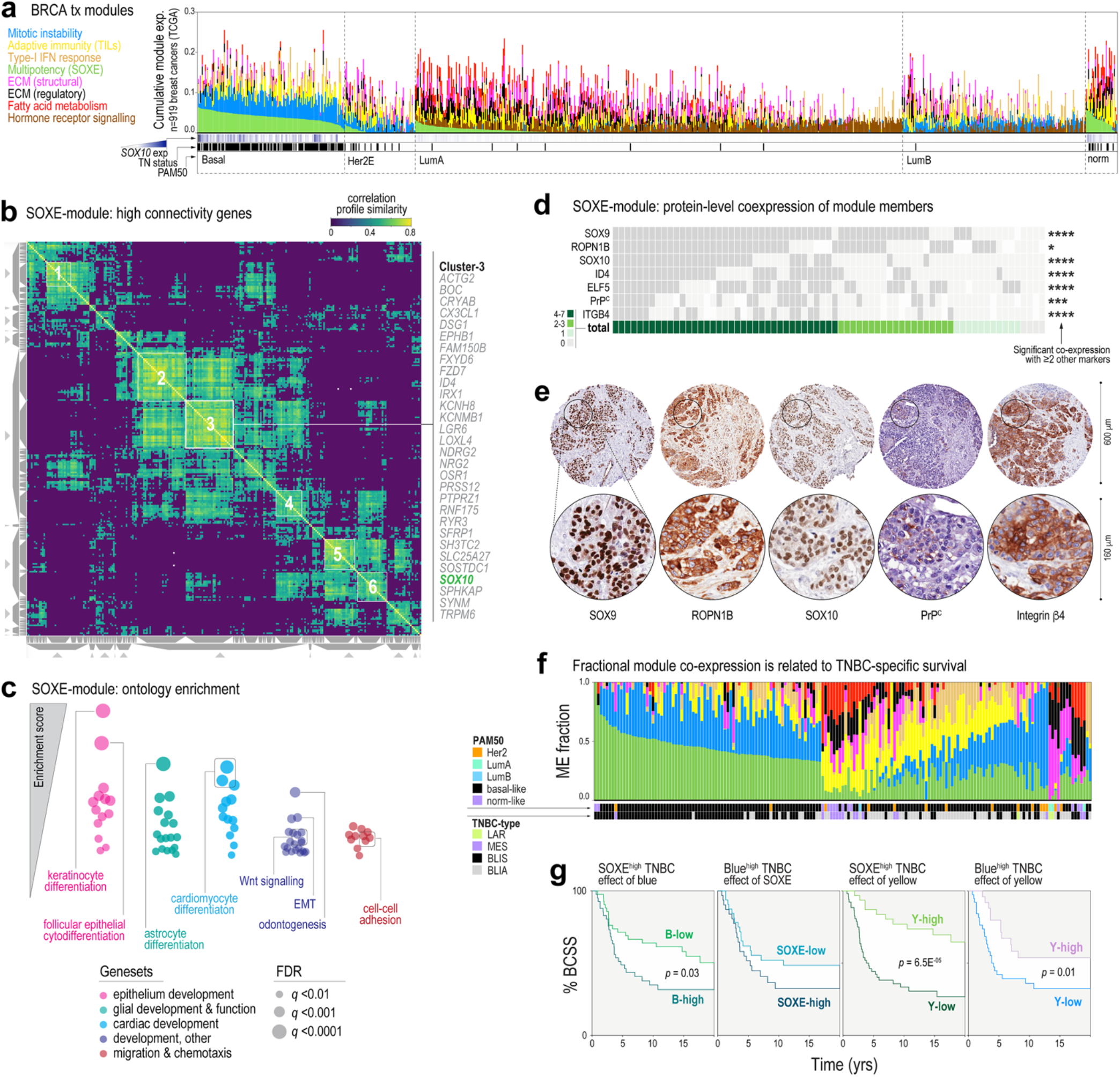
SOX10’s regulatory network is associated with multipotency, cell migration and poor prognosis in TNBC. (**a**) Relative expression of eight predominant transcription modules in human breast tumours, according to PAM50 subtype (TCGA dataset). (**b**) SOXE-module co-expression profile similarity matrix, clustered to highlight genes with very highly coordinated expression. Similarity is based on cosine-distance and has a maximum value of 1. *SOX10* mapped to one of six module sub-clusters, the members of which are shown to the right of the matrix. See also Figs S4a-b. (**c**) Summary of results from unsupervised geneset enrichment analysis of the breast cancer transcriptome after ordering transcripts according to their correlations with SOXE-module expression (denoted by the ME value, TCGA dataset). (**d**) Tile plot showing overlapping expression of SOXE-module representatives. For each protein, significant co-expression with ≥2 other module members is indicated by a Fisher’s Exact test result (\**p*<0.05; \*\*\**p*<0.001; \*\*\*\**p*<0.0001). Refer to methods (Table-M1) for scoring criteria. (**e**) IHC staining of representative SOXE-module nodes in serial sections from the same tumour. (**f**) Proportional expression of all eight tx modules (coloured as for (a)) in TNBCs annotated with PAM50 and TNBC subtypes (METABRIC dataset; *LAR, luminal androgen receptor-like; MES, mesenchymal; BLIS, basal-like immune-suppressed; BLIA, basal-like immune-activated^39^).* (**g**) Kaplan Meier analysis of METABRIC TNBCs expressing different proportions of the three predominant TNBC modules. BCSS, breast cancer-specific survival. ME fraction thresholds for classifying cases as high or low were 0.33 for SOXE/blue and 0.1 for yellow.

**Table-1.**
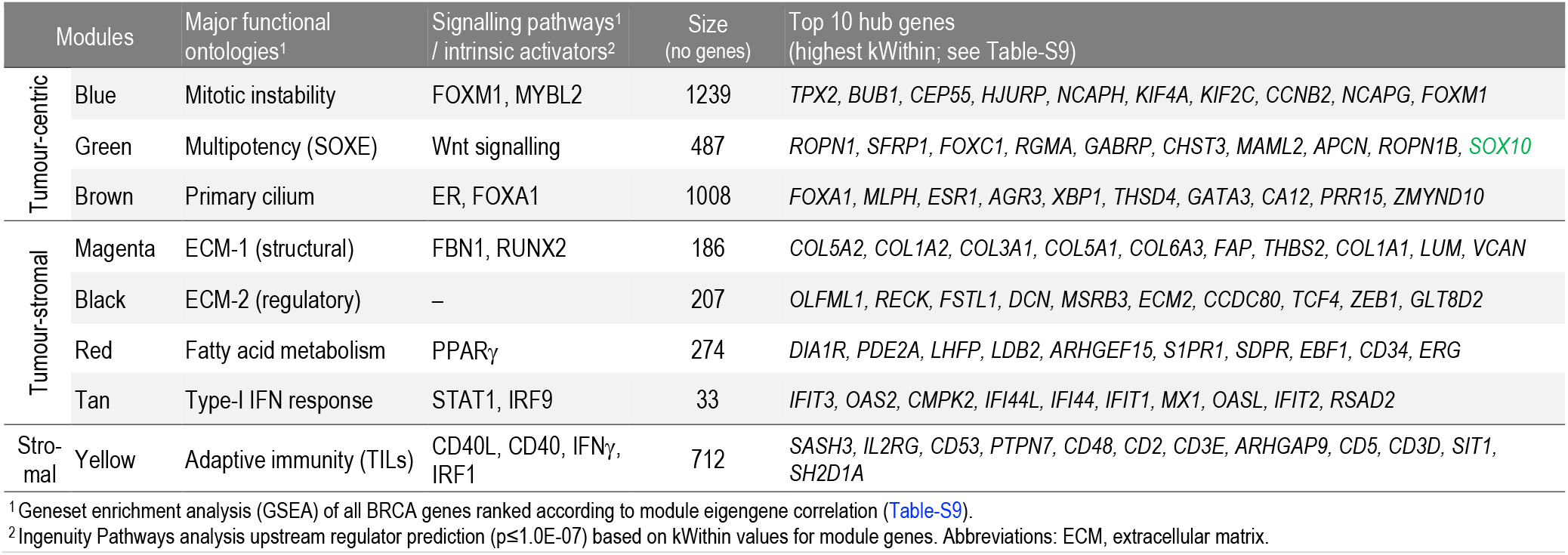
Key features of eight breast cancer transcription modules extracted by WGCNA.

IHC analysis of six other module members confirmed that their co-expression in TNBC holds true at the protein level (Fig-3d), with staining often observed in the same cells within individual tumour-rich tissue cores (Fig-3e). Consistent with the defining features of TNBCs – de-differentiation, genomic instability, high mitotic index and the presence of TILs – TNBCs express variable proportions of primarily three modules: green (SOXE), blue (mitotic instability) and yellow (TILs) (Fig-3f). Kaplan-Meier analysis showed that cases expressing high levels of both SOXE and mitotic instability modules had shorter survival compared to those with predominant expression of one or the other, while co-expression of the yellow module was associated with better prognosis, consistent with the protective effect of TILs in TNBC^54^ (Fig-3g, Fig-S4d).

### The SOXE-module represents the shift from a luminal progenitor to a NCSC-like state

Ontology analysis showed that the SOXE-module includes genes typically expressed in differentiating glia, cardiomyocytes and odontoblasts, which all descend from NCSCs. In fact, developmental genes comprised a large proportion of SOXE-module hubs (genes with the highest network connectivity and centrality values; Fig-4a; Table-S10), hence represent points of maximal module vulnerability. These include cell fate regulators *ELF5, FOXC1* and *SOX10;* Wnt/ß-catenin signalling genes *SFRP1, MAML2* and *TRIM29;* and embryonic cell migration and neuronal development genes *RGMA, ROPN1, ROPN1B, MID1* and *APCN.*

**Figure-4.**
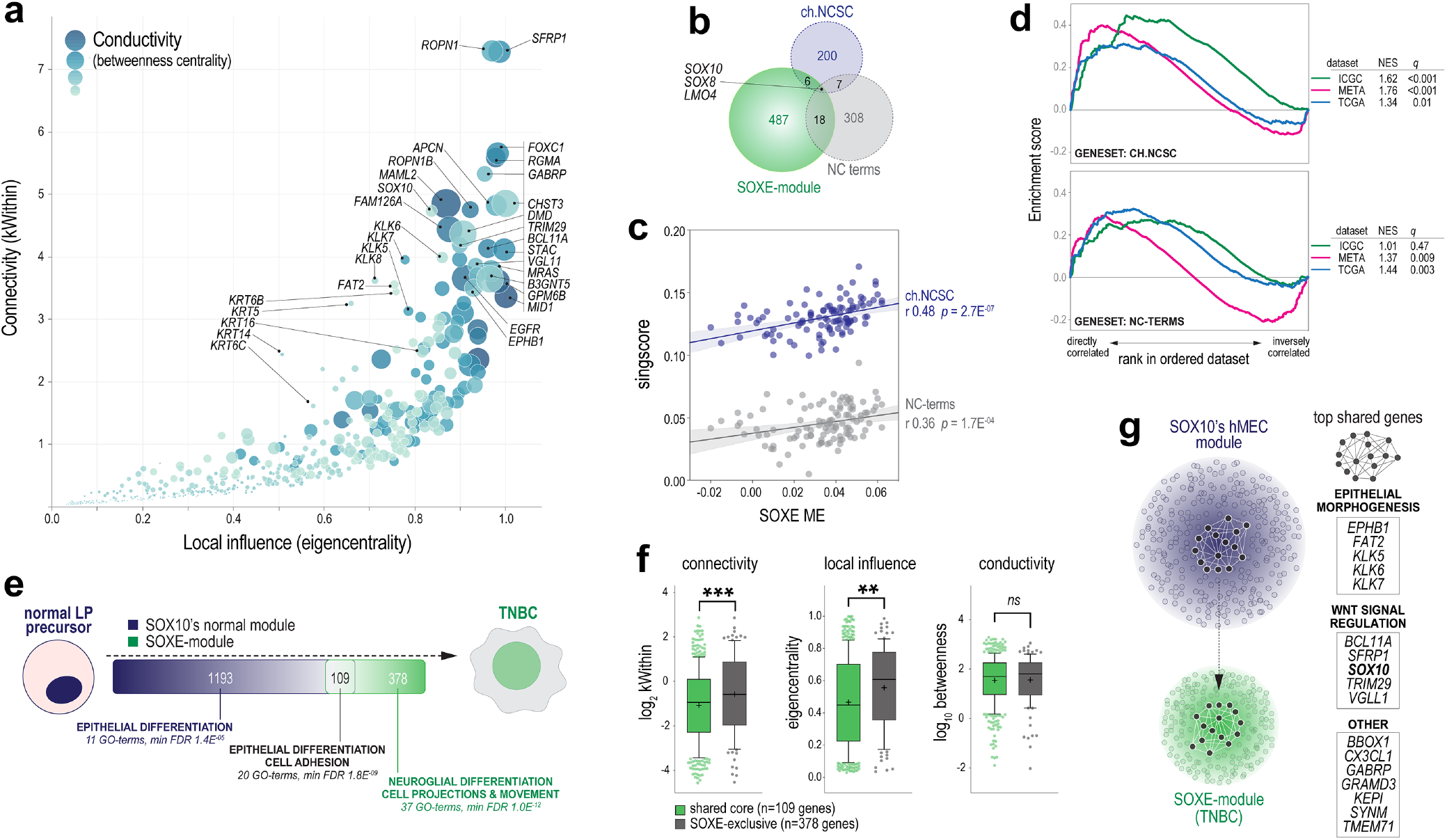
The SOXE-module drives the transition from normal mammary epithelial stem/progenitor to NCSC-like phenotypic states. (**a**) Relative importance of SOXE-module genes in terms of influence over network architecture and information flow. High values across the following three network metrics are indicative of ‘hubness’: *kWithin:* intramodular ‘connectivity’ based on weighted correlations with all other module genes; *Eigencentrality:* considers the connectivity of each node’s nearest neighbours as an indicator of ‘local influence’; *Betweenness centrality:* measures ‘conductivity’ based on each node’s position along the shortest paths between other nodes (genes with high betweenness are information conduits that lie on many shortest-paths between nodes. Key hub genes are indicated (see Table-S10 for the full dataset). (**b**) Chick (ch.)NCSC and neural crest (NC) terms genesets used for phenotypic analysis are largely independent from each other and from the SOXE-module. (**c**) Correlations between SOXE ME values and NCSC genesets *(singscore* expression values) in TNBC (n=106 TCGA cases with tumour cellularity ≥0.6). Correlation coefficients (r) and *p*-values shown. (**d**) GSEA using three TNBC gene expression datasets (ICGC, METABRIC, TCGA). Normalised enrichment scores (NES) and corrected *p*-values *(q)* shown. (**e**) Overlap between members of the SOXE-module and SOX10’s normal breast module (from *de novo* module identification on n=97 TCGA normal breast samples; Table-S12). Generic ontology enrichment results are summarised (full GO term lists in Table-S13). (**f**) Comparison of network structure and information flow metrics (as for (a)) between shared and SOXE-module-exclusive genes. Groups were compared using Mann-Whitney tests (\*\**p*=2.4E^-03^; \*\*\**p*=5.6E^-04^). (**g**) Model depicting the mammary epithelial progenitor gene regulatory network core being sustained through transformation and rewired as the SOXE-module in TNBC. Shared hub genes are listed.

To directly investigate if the SOXE-module is associated with NCSC phenotypic mimicry, as has been reported for Sox10 in mouse mammary tumour cells^24^, we performed expression and enrichment analyses using two independent genesets: (1) 308 genes represented in at least two of the 78 terms matching ‘neural crest’ in the gene ontology database (‘NC terms’); and (2) transcripts specific to migratory, Sox10+ NCSCs in chick embryos (‘ch.NCSC’; n=200 genes)^55^, representing Sox10’s most primitive transcription program (Table-S11). Except for *SOX10, SOX8* and *LMO4,* there is minimal overlap between the SOXE-module and these genesets (Fig-4b), but their expression is strongly correlated (Fig-4c). This was confirmed by geneset enrichment analysis (GSEA; Fig-4d). Hence, the SOXE-module confers transcriptomic similarity to NCSCs.

Since several SOXE-module genes (e.g., *SOX10, SOX9, LGR6, ELF5)* are key regulators of normal hMEC states^56^, we hypothesised that the SOXE-module might evolve from the deregulation of a lineage differentiation program expressed in TNBC’s normal cellular precursors. Module preservation analysis using RNAseq data from TCGA normal breast samples indicated that the SOXE-module does not exist as an interconnected unit in the normal breast transcriptome (Fig-S4e). But after performing *de novo* WGCNA module identification on this dataset (Table-S12), we found that *SOX10’s* normal breast module overlaps with the TNBC-specific SOXE-module significantly more than expected by chance (Fig-4e; 109 shared genes, Chi Square *p*=2.8E^-26^).

Both ‘normal-exclusive’ and ‘shared’ genes were enriched with epithelial differentiation ontologies, with cell adhesion distinctly over-represented in the shared set (Fig-4e, Table-S13). According to network influence metrics, the shared genes were significantly more important to the SOXE-module than SOXE-exclusive genes (Fig-4f, Fig-S4f). These data suggest that while SOXE-exclusive genes are primarily responsible for conferring NCSC-like attributes, genes ‘inherited’ from TNBC’s normal precursors are comparatively more important to the SOXE-module’s regulatory structure. Together, these data suggest that SOXE-module and its associated NCSC-like phenotype arises because a core set of epithelial differentiation and adhesion genes becomes rewired during TNBC development (Fig-4g).

### Genomic and epigenomic determinants of the NCSC-like transcriptional shift in TNBC

To address the central question of what drives this transcriptomic shift, we analysed case-matched gene copy-number (CN), RNAseq and WGCNA data (TCGA cases). Candidate module drivers were defined as those for which both CN and expression correlated significantly with SOXE-ME values. 182 genes met these criteria (130 gains, 52 losses), of which 140 (77%) are part of large chromosomal alterations: 6p21-22 (gained/amplified in 56.7% of TNBC cases), 8q22-24 (gained/amplified in 78.7%), 9q34 (lost in 59.6%) (Fig-S5a). SOXE-module genes were over-represented amongst the positively correlated genes (25/130 (19.2%) and had increased CN and expression in SOXE^high^ TNBC; ChiSq *p*=9.7E^-31^; Fig-5a). However, network influence metrics for these 25 were no higher than other module genes (Fig-5b). Hence, the SOXE-module may be augmented by increased CN of some of its component genes, but this seemed unlikely to be an early or dominant driver of module evolution.

**Figure-5.**
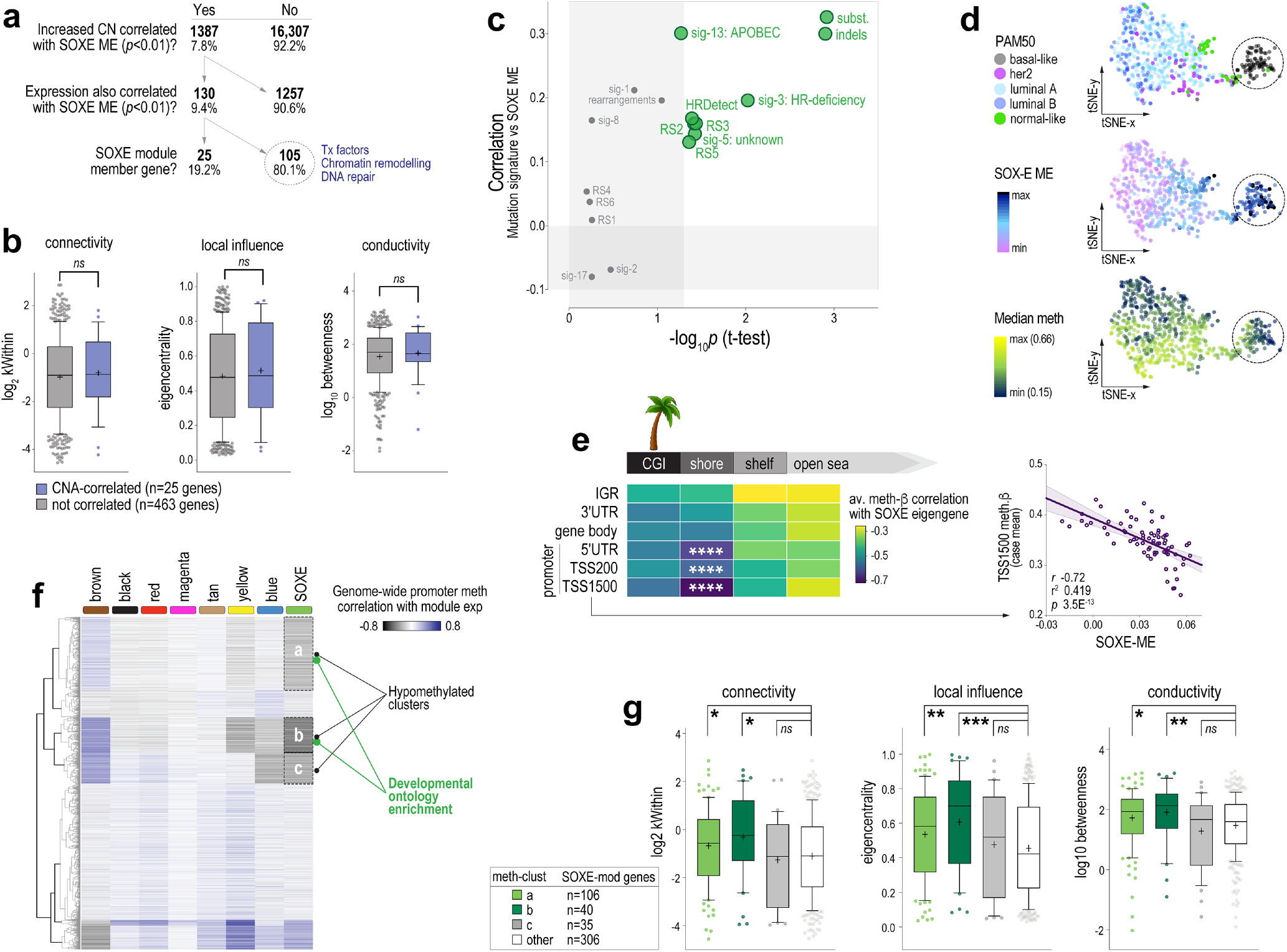
The SOXE-module is driven by erosion of lineage-specific epigenetic marks. (**a**) Decision tree for identifying candidate copy-number alteration (CNA) drivers of the SOXE-module. Of 17,694 genes with case-matched GISTIC, RNAseq and WGCNA data, CN and expression of 130 correlated with the SOXE-module in TNBC, including 25 SOXE-module nodes. (**b**) Network influence metrics for SOXE-module nodes coloured according to candidate CN driver status (intramodular connectivity (kWithin), local influence (eigen-centrality), and conductivity (betweenness centrality) defined in the caption for Fig-4a). No significant differences by ordinary ANOVA test. (**c**) Relationship between SOXE-module levels and mutation signatures in ICGC TNBCs (COSMIC v2 SigProfiler and HRDetect on n=74 ICGC TNBCs)^45^. Associations are depicted according to the correlation between SOXE-ME values and signature event count (y-axis); and by the significance of average SOXE-ME differences between ICGC TNBCs with low (quartile-1) vs higher (quartile 2-4) signature burden. (**d**) t-Distributed Stochastic Neighbour Embedding (t-SNE) visualisation of genome methylation profile similarities amongst cases in the BRCA-TCGA 450k methylation array dataset. Panels are coloured according to PAM50 intrinsic subtype, SOXE-ME values or global median methylation-ß values. Circled cases are epigenetically divergent, basal-like TNBCs that express high levels of the SOXE-module and have eroded methylomes. (**e**) Correlation analysis summary showing relationships between SOXE-ME values and region-specific methylation (n=75 TCGA TNBCs, tumour cellularity ≥0.6; n=215,323 probes after quality filtering); \*\*\*\**p*<1.0E^-07^. *CGI, CpG island; IGR, intergenic region; TSS, transcription start site; UTR, untranslated region. Solo-WCpGW: consensus sequence for late replicating loci demethylated via replicative senescence.* (**f**) Unsupervised clustering of the BRCA-TCGA 450k methylation dataset according to ME correlation. Data shown are minimum correlation coefficients of ME values versus gene-averaged methylation beta data from promoter region probes (TSS1500, TSS200, 5’UTR). Of three clusters inversely correlated with SOXE-module expression, two (a,b) were enriched with developmental ontologies (Table-S14). (**g**) Network influence metrics for SOXE-module genes in the hypomethylated clusters versus other SOXE-module genes, as for (b). Ordinary ANOVA *p*-values: \**p*<0.05; \*\**p*<0.01; \*\*\**p*<0.001; *ns*, not significant.

Next, we investigated whether mutational processes that shape the breast cancer genome could be involved. To this end, we utilised mutational signature and WGCNA data for matching ICGC cohort cases^45,57^. There were direct relationships between the SOXE-module and overall mutation burden (substitutions and small insertion-deletions (indels)), as well as specific signatures of genome instability (rearrangement sigs (RS)3 and RS5), homologous recombination (HR)-directed repair of double-strand DNA breaks (DSBs) and genome editing (sig-3: HR deficiency; HRDetect; sig13: APOBEC; Fig-5c).

APOBEC activity and DSB repair are both indirectly demethylating. For example, 5-methyl cytosine (5mC) loss occurs as a result of APOBEC-mediated genome editing and/or during the repair of edited bases, and DSB repair has been causally linked to the progressive loss of 5mC during cellular ageing^58,59^. Therefore, we hypothesised that evolution of the SOXE-module in TNBC may be related to epigenetic dysregulation. Consistent with this idea, the 105 CN-driven SOXE-module correlates (i.e., those not part of the SOXE-module itself; Fig-5a) were enriched with transcription factor, chromatin remodelling and DNA repair genes (Fisher’s Exact *p*<0.001). Furthermore, visualising SOXE-module expression relative to overall methylome profile using tSNE maps showed that SOXE-ME values were highest in the most epigenetically divergent tumours (Fig-5d).

To investigate this further, we then correlated SOXE-ME values with probe-level methylation data directly, in the following regional categories: CpG islands (CGIs), CGI shores, shelves or open sea regions at transcription start site (TSS) regions, untranslated regions (UTRs), gene bodies or intergenic regions (IGRs). We also quantified methylation at ‘solo-WCpGW’ sites at late-replicating, heterochromatic loci, which act as a biomarker of replicative senescence^60^ and are hypomethylated in breast tumours compared to hMECs (Fig-S5b). There was actually no relationship with solo-WCpGW sites (Fig-S5c); but there was a striking inverse correlation between SOXE-ME values and genome-wide promoter methylation; particularly at CGI shores, the substrate for lineage-specific methylation in adult tissues (Fig-5e, Fig-S5c). These data indicate that SOXE-module expression and connectivity are directly proportional to promoter demethylation in TNBC (Fig-5e). There was no such relationship with any other module in TNBC (Fig-S5d).

Having established that SOXE-module levels correspond with erosion of tissue-specific 5mC marks, we then built a correlation matrix from ME and genome-wide promoter methylation data (TCGA) and performed unsupervised clustering to look for evidence of epigenetic control. Indeed, the SOXE-module had a distinct promoter methylation signature, including three clusters of genes that are hypomethylated when SOXE-module strength is highest, of which two were enriched with developmental ontologies (Fig-5f, Table-S14). Only 10% of these genes correspond to SOXE-module genes, but this 10% is enriched with influential hub genes (Fig-5g), suggesting a higher level of epigenetic control over module structure and information flow. We then used GSEA to test enrichment of the SOXE-associated promoter methylome with NCSC genesets. Like the transcriptome (Fig-4d), the methylation landscape associated with SOXE-module expression was also enriched with NCSC genes (NC terms: normalised enrichment score (NES) −1.5; *q*=6.0E^-03^; Ch.NCSC: NES −1.3; *q*=3.6E^-02^).

Finally, we investigated direct demethylation processes as potential enablers of SOXE-module formation by cross-referencing SOXE-ME values from our three WGCNA datasets (TCGA, ICGC, METABRIC) against the expression of demethylases in the EpiFactors database^61^. There were direct associations with APOBEC3A/3B cytosine deaminases and *TET1* (Fig-S5e). TET dioxygenase enzymes catalyse the first step of 5mC demethylation and are involved in processes requiring cell states to be reset or adjusted, such as methylome erasure in preimplantation embryos, and epigenetic plasticity in brain regions that facilitate learning and memory. *TET1* is a maintenance demethylase that prevents methylation spreading from silenced loci, particularly at CGI shores^62,63^. It has been causally implicated in TNBC metastasis^64^ and our findings suggest this may be at least partly due to reinforcement of the SOXE-module.

In summary, these analyses show that the SOXE-module correlates with APOBEC activity, DSB repair and *TET1* expression, which are all associated with genome-wide demethylation. Given that the SOXE-module’s dominance over the TNBC transcriptome is proportional to demethylation at promoter CGI shores, we postulate that the demand for DSB repair drives progressive methylome erosion during TNBC development, de-programming LP cells while making them vulnerable to reprogramming by NCSC fate specifiers like SOX10 (Fig-6).

**Figure-6.**
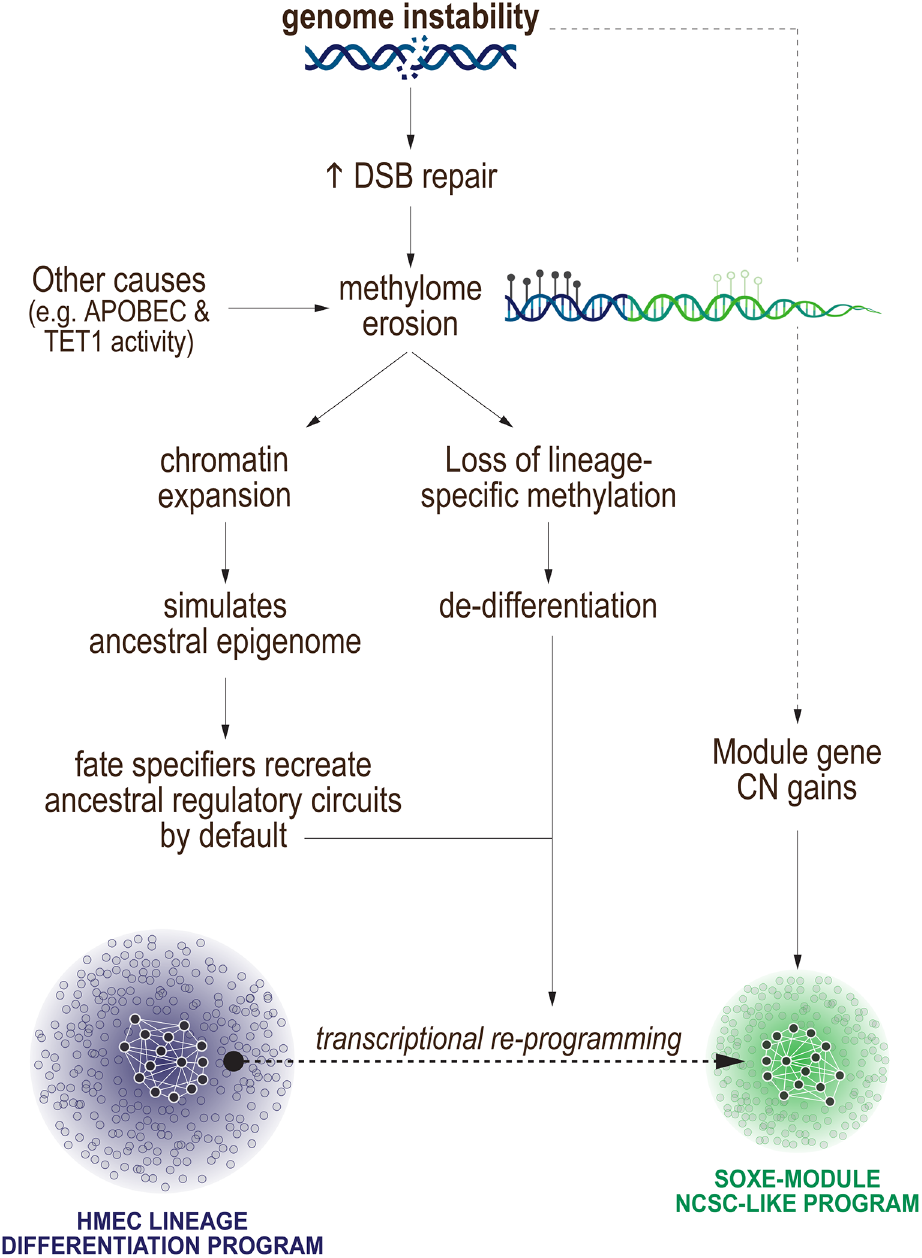
Proposed contribution of genome instability and epigenome erosion to transcriptional reprogramming and NCSC-like phenotypic mimicry during TNBC development.

### Other SOX10+ cancers with eroded epigenomes exhibit NCSC phenotypic mimicry

Genome-wide hypomethylation is also prevalent in other cancers, particularly those of high histologic grade^65,66^. If this simulates to some extent the expanded chromatin landscape of NCSCs (Fig-7a), we reasoned that other epigenetically eroded, SOX10+ cancers may also exhibit NCSC phenotypic mimicry. All known SOX10+ malignancies arise in tissues maintained by SOX10+ resident stem cells, thus SOX10 could have an early and enduring influence on cell state reprogramming during the progression of these cancers (Fig-7b). We investigated this idea by performing systems-level analysis of TCGA RNAseq data from 22 solid tumour types.

**Figure-7.**
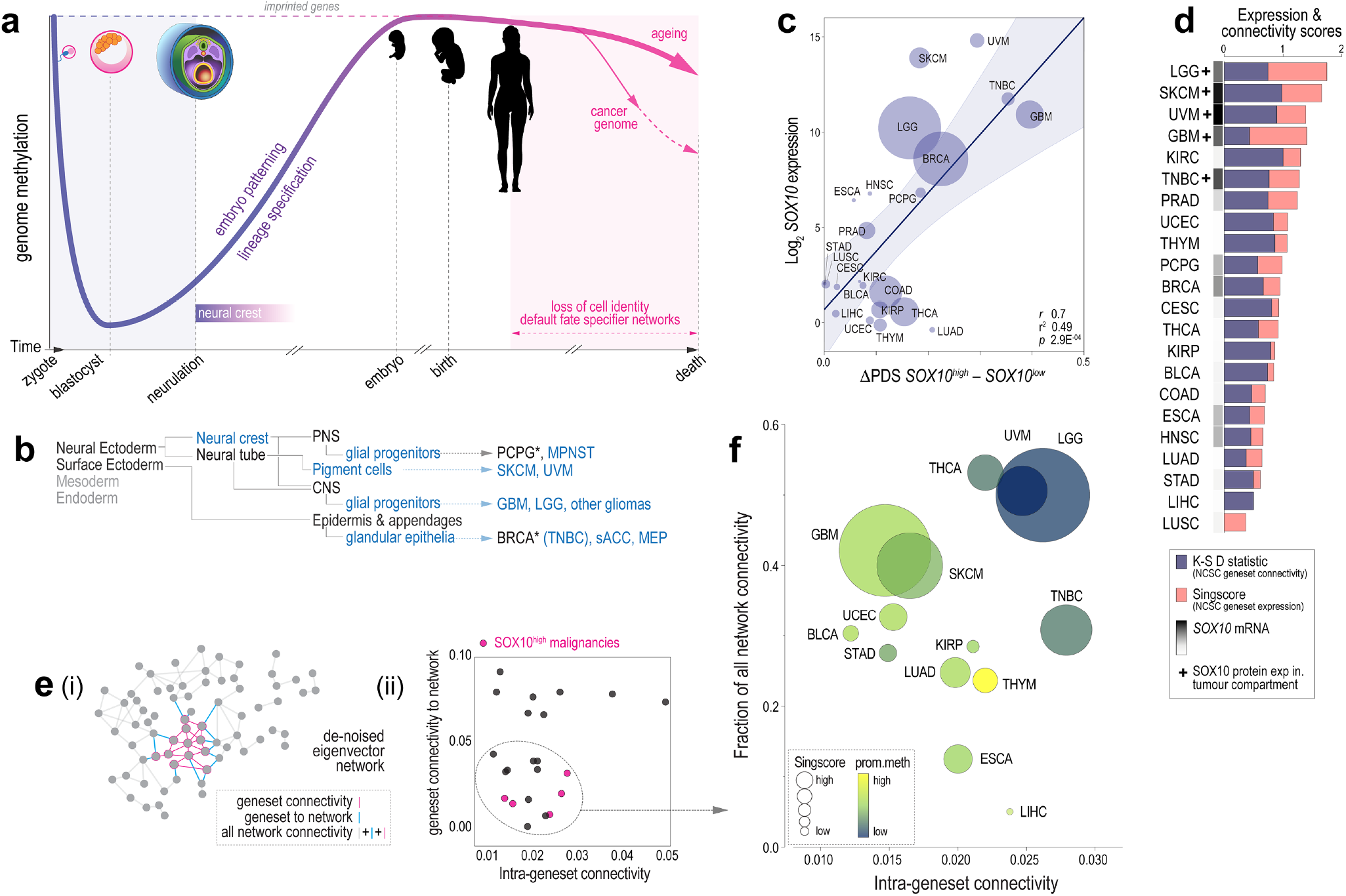
SOX10+ cancers have eroded methylomes and transcriptomes dominated by a NCSC-like gene regulatory network. (**a**) Changes in genome methylation over time. Neural crest is the earliest developmental state to express *SOX10* and exists when the embryonic genome is broadly hypomethylated and expanded. Cell states are defined by methylation barcoding in differentiating cell types. These marks are progressively eroded during normal cellular ageing; accelerated in cancer because of replicative erosion, mutational and DNA repair processes. (**b**) *SOX10* expression traced from malignancy back through the normal cellular precursors and developmental lineages (blue text = SOX10+). Adapted from LifeMap^70^. *\*SOX10* RNA detected in PCPG and nonTN BRCA samples does not originate from the tumour compartment (sustentacular support cells for PCPG, and contaminating normal mammary epithelia for nonTN BRCA^71^). (**c**) Pathifier analysis of ch.NCSC and NC-terms genesets in *SOX10^high^* versus *SOX10^low^* counterparts of 22 cancer types. Data shown are the mean differences in pathway dysregulation scores (DPDS) of the two genesets between high and low groups of each histology, in relation to baseline *SOX10* RNA expression (TCGA RNAseq data), and the statistical significance of DPDS for each malignancy proportional to circle size (2-tailed t-tests). There was a strong relationship between *SOX10* expression and DPDS (Pearson correlation coefficient (r), goodness of fit (r^2^) and significance (p) shown). (**d**) NCSC geneset expression (Singscore) and connectivity (Kolmogorov-Smirnov D statistic) across cancers compared to *SOX10* expression. (**e**) Pan-cancer analysis of ch.NCSC geneset connectivity after network de-noising. (i) Diagram defining key network metrics. (ii) Geneset-network (Y-axis) and intra-network (X-axis) connectivity for 22 cancer types. (**f**) ch.NCSC geneset connectivity (X-axis) versus the fraction of all connectivity attributed to the geneset after de-noising. Singscore geneset expression and global promoter methylation levels are superimposed. Abbreviations: BLCA, Bladder Urothelial Carcinoma; BRCA, Breast invasive carcinoma (non-TN cases); CESC, Cervical squamous cell carcinoma and endocervical adenocarcinoma; COAD, Colon adenocarcinoma; ESCA, Oesophageal carcinoma; GBM, Glioblastoma multiforme; HNSC, Head and Neck squamous cell carcinoma; KIRC, Kidney renal clear cell carcinoma; KIRP, Kidney renal papillary cell carcinoma; LGG, Brain Lower Grade Glioma; LIHC, Liver hepatocellular carcinoma; LUAD, Lung adenocarcinoma; LUSC, Lung squamous cell carcinoma; MEP, myoepithelioma; MPNST, malignant peripheral nerve sheath tumour; PCPG, Pheochromocytoma and Paraganglioma; PNS, peripheral nervous system; PRAD, Prostate adenocarcinoma; sACC, salivary adenoid cystic carcinoma; SKCM, Skin Cutaneous Melanoma; STAD, Stomach adenocarcinoma; THCA, Thyroid carcinoma; THYM, Thymoma; UCEC, Uterine Corpus Endometrial Carcinoma; UVM, Uveal Melanoma.

First, we characterised NCSC geneset expression using the *singscore* tool^67^, which revealed a range of expression levels that generally corresponded with that of *SOX10* (Fig-S6a). We then analysed SOX10’s influence over NCSC gene expression using *Pathifier,* which provides a ‘pathway de-regulation score (PDS) representing the collective test sample variance around a principal curve trained on controls^68,69^. After stratifying each cancer cohort into two subgroups based on median *SOX10* mRNA level, we trained *Pathifier* on NCSC geneset expression in *SOX10^low^* (control) samples, then quantified the degree of change in corresponding *SOX10^high^* (test) samples. This revealed a direct relationship that was most pronounced for *SOX10^high^* cancers – melanoma (UVM, SKCM), glioma (GBM, LGG) and TNBC (Fig-7c). Hence, expression of *SOX10* in cancer is associated with expression of other NCSC genes.

As the NCSC genes could be involved in a range of distinct processes in adult tissues, we also performed network analysis to infer co-functionality amongst these genes with more confidence. Specifically, we asked whether NCSC genes are more connected than expected by chance, given the unique transcriptome network structure of each malignancy (Fig-S6b). After building correlation matrices from the 22 RNAseq datasets, we iteratively quantified the median connection strength amongst sets of 60 genes at a time, selected either at random from the NCSC genesets, or the entire transcriptome as a control (1000 permutations). NCSC geneset connectivity was indeed above background in all cancers (lower median inter-transcript distance compared to the control; Figs S6c-d), indicating a degree of coordinated regulation. The ch.NCSC geneset was significantly more connected than the NC terms list (Fig-S6e), possibly because the former was identified on the basis of co-expression^55^, while the NC terms set was simply compiled from GO database entries.

The pan-cancer analyses indicated that SOX10-expressing malignancies not only express higher levels of NCSC genes, but also that fluctuations in the expression of these genes across large tumour sample cohorts tend to be coordinated (Fig-7d). However, there was still a large amount of ‘noise’ in the correlation matrices, with large variation in the distribution of random permutation data for some cancers and not others (Fig-S6f). To correct for these structural differences, we took a novel, computationally intensive approach to de-noise the cancer-specific correlation matrices; transforming them into ‘concurrence matrices’ that are stripped of network-wide influences and noise eigenvalues. This drastically pruned the original networks, and yet NCSC geneset connectivity signals remained detectable. We then derived the following metrics for each malignancy: (1) median eigenvectors of all ch.NCSC gene pairs (intra-geneset connectivity); and (2) median eigenvectors of *any* gene pair involving ch.NCSC genes, reflecting connectivity to all other genes (geneset-network connectivity) (Fig-7e(i)). Interestingly, the denoised networks loosely fell into two clusters, with lower values indicating geneset coordination that is independent from the rest of the network (Fig-7e(ii)).

For the lower cluster, we calculated intra-geneset connectivity as a fraction of all connectivity remaining after the denoising process. This showed that NCSC genes are not only strongly interconnected in SOX10*+* cancers, but account for a substantial fraction of all true (non-noise) eigenvalues, indicating the NCSC program is a dominant influence within the transcriptomes of these cancers. Superimposing *singscore* and genome-wide promoter methylation data further consolidated the positive association between epigenome erosion and coordinated expression of NCSC genes in SOX10+ cancers (Fig-7f).

## Discussion

Heterogeneity has emerged as a major bottleneck to effective sub-classification and treatment of cancer, and TNBC is no exception. Post-treatment relapse occurs through clonal expansion of cells with pre-existing, advantageous mutations, but also cell state changes brought about by adaptive epigenetic remodelling – a phenomenon that unites the ‘cancer stem cell’ and ‘epigenetic progenitor’ models of cancer^72^. The intrinsic plasticity of TNBC is problematic because existing therapies cannot eradicate a shifting target. Early evidence implies that blocking this capability with epigenetic therapy may improve treatment efficacy, but this will require a deeper understanding of how phenotypic plasticity evolves^73^. TNBC exhibits genome-wide hypomethylation, which evidently drives de-differentiation by destroying the state-defining epigenetic barcode of its normal cellular precursor, the LP cell^14–17,72,74^. Differential methylation at certain genomic loci is prognostic in TNBC^22^, and myriad studies have helped to decipher the mechanistic contributions of individual writers, readers and erasers of epigenetic marks, but the phenotypic manifestations of genome-wide 5mC loss have not been extensively studied.

Consistent with evidence that cell state specifiers are important for phenotypic plasticity, including functional analysis of Sox10 in experimental mice^29,32^, our systems-level network studies show that *SOX10’s* TNBC-specific regulatory module confers transcriptomic similarity to highly plastic NCSCs. We traced a cluster of super-connected SOXE-module genes back to tissue-resident mammary stem and progenitor cells. In contrast to normal breast where it was associated with epithelial lineage differentiation, in TNBC this core was connected to Wnt signalling, neuroglial differentiation and embryo patterning genes. Apart from an ontology enrichment profile similar to NCSCs, the SOXE-module also conferred transcriptomic similarity to the Sox10+ migratory NCSC compartment from chick embryos^55^. We also identified SOXE-module hub genes as points of maximum network vulnerability as candidate therapeutic targets. In support of this approach, two of these – *BBOX1* and *BCL11A* – have already been validated as such in TNBC^75–79^.

In terms of understanding SOXE-module evolution, we focused on established drivers of TNBC development – genomic instability, large-scale CNAs and defective DNA repair. We identified several processes that correlate significantly with the SOXE-module eigengene (DSB repair, APOBEC and TET1 activity, which are all demethylating); but most discernibly, the loss of lineage-specific methylation marks at CGI shores. This suggests that NCSC-like re-programming occurs concomitantly with epithelial deprogramming in TNBC and is consistent with tracing analyses (Fig-4) that mapped part of the SOXE-module core in the epithelial lineage differentiation module of normal breast samples.

A direct link between DSB repair and epigenome erosion was established only very recently^58,59^. These compelling studies demonstrated how over time, DSB repair drives epigenome ‘smoothening’ and a loss of cellular identity during ageing. The implications of this in cancer biology have not yet been realised but will likely be far-reaching, given the heightened demand for DSB repair in various malignancies. Other mechanisms could also contribute to methylome erosion in cancer, such as reduced availability of 5mC substrates as can occur with metabolic reprogramming^80^. Accepting that there are multiple causes, epigenetic erosion is nonetheless prevalent in poorly differentiated cancers^65,66^. We found that promoter hypomethylation also correlates with the NCSC-like phenotype in other cancers that express SOX10. The one exception was thyroid cancer, which does not express SOX10^81^, but does express other NCSC specifiers (*SOX9*, *ETS1, TFAP2, RXRG*^82–85^); and interestingly, thyroid hormone is an obligate requirement for NCSC migration during embryogenesis^86^. The self-reinforcing gene regulatory networks that operate in NCSCs are amongst the most evolutionarily conserved in vertebrates^25,87^. Therefore, we postulate that when the broadly expanded chromatin landscape of the early embryo is simulated in epigenetically eroded cancers, fate specifiers like SOX10 may recreate their ancestral regulatory circuits by default.

Links between development and cancer were first documented in the 19^th^ Century^88,89^. Key milestones in the study of developmental phenotypic mimicry include the discovery that embryonic markers are ‘reexpressed’ in malignant tissue; that melanoma cells partially ‘hijack’ the NCSC transcriptome in order to metastasise^90^; and that embryonic stem cell gene expression signatures are prevalent in poorly differentiated tumours (including basal-like breast cancer)^91^. But for many years the mechanistic basis of this phenotypic mimicry has remained opaque.

Finding that both SOX10 and its cancer-specific regulatory module are poor prognostic indicators in TNBC, we mined high-dimensional tumour (epi)genomic datasets to address the question of *how* developmental phenotypes evolve during cancer development; an approach only possible because of international cancer (epi)genome sequencing efforts^43–45^. One limitation of this approach is that tumour datasets suffer from considerable stochasticity and noise. To mitigate this, we used data from orthogonal platforms, applied tumour cellularity cut-offs where appropriate and cross-referenced WGCNA data against established biological and clinicopathologic markers. Crucially, we also developed a novel network eigenvalue ‘denoising’ pipeline, based on preserving only those eigenpairs whose eigenvalues are outside the expected relevant random matrix eigenspectrum.

## Conclusions

Our data indicate that the extent of genome-wide promoter methylation loss in SOX10+ malignancies correlates with their transcriptomic similarity to NCSCs – the earliest developmental cell state programmed by SOX10 activity and one synonymous with migration, multipotency and phenotypic plasticity. Concerning TNBC development, we propose that progressive erosion of the LP-specific epigenome drives dedifferentiation while simultaneously making cells vulnerable to NCSC-like reprogramming. Broadly, these findings support preclinical data^19^’^23^ on the potential for chromatin remodelling inhibitors to limit the development of chemoresistance.

## Declarations

### Ethics approval and consent to participate

Human research ethics approval was obtained from the Royal Brisbane and Women’s Hospital (2005000785), The University of Queensland (HREC/2005/022) and North West Greater Manchester Central Health (15/NW/0685). Written patient consent to use tissue for research purposes was obtained where required under the conditions of these approvals and all samples were de-identified in the analytical database. This study complies with the World Medical Association declaration of Helsinki.

### Consent for publication

Not applicable

### Availability of data and materials

All data generated or analysed during this study are included or referenced in supplementary information.

### Authors’ contributions

Conception & design: JMS, XMDL, KN, AR, DVN, PTS, SRL. Data collection/contribution: JMS, XMDL, KN, AR, AH, AEMR, ML, ACV, JRK, AJD, MM, EK, PKdC, IG, FA, JMWG, CO, KKK, JB, GCT, ATG, EAR, IOE, DVN, PTS. Data analysis: JMS, XMDL, KN, AR, AH, SL, DVN. Manuscript drafting: JMS, AEMR, DVN, PTS, SRL. All authors read and approved the final manuscript.

## Acknowledgements

We thank the many thousands of patients who have donated tissue for cancer research, and clinical staff who facilitate biobanking, particularly the Brisbane Breast Bank and Pathology Queensland. We acknowledge the support of Metro North Hospital and Health Services for the collection of the clinical subject data and clinical Subject materials. We are grateful to Dr Lynne Reid and Clay Winterford for valuable contributions to histological analyses; Dr Katia Nones (QIMR Berghofer) who supervised XMDL; Dr Chris Schmidt (QIMR Berghofer) and A/Prof Alex Swarbrick (Garvan Institute) for donating cell lines; Dr William Cockburn and clinical staff (Wesley Hospital) for normal breast tissue collections; Dr Nic Waddell and Olga Kondrashova (QIMR Berghofer) for supportive data analyses; and Drs Juliet French (QIMR Berghofer) and Delphine Merino (Olivia Newton John Cancer Research Institute) for critical feedback. This study makes use of data generated by the Molecular Taxonomy of Breast Cancer International Consortium, funded Cancer Research UK and the British Columbia Cancer Agency Branch.

## Supplementary information

**Supplementary file 1**: Materials, methods & data availability.

**Table-M1**: Antibodies and immuno-staining conditions used in this study.

**Table-M2:** Module membership correlation between TCGA and METABRIC datasets.

**Figure-M1:** Breast cancer WGCNA network topological overlap.

**Figure-M2**: Module preservation.

**Figure-M3:** Network schematic defining nodes, hubs, modules and node influence metrics.

**Supplementary file 2**: Figures.

**Figure-S1**: Data supporting Fig-1

**Figure-S2**: Data supporting Fig-2

**Figure-S3:** Gene ontology profiles of SOX10-high and -low TNBCs

**Figure-S4**: Data supporting Fig-3

**Figure-S5**: Data supporting Fig-5

**Figure-S6**: Data supporting Fig-7

**Supplementary file 3**: Tables.

**Table-S1**. Reagent & resources table (methods supplement).

**Table-S2**. Association of SOX10 expression with histopathological parameters in breast cancer, and results of multivariate regression analysis.

**Table-S3**. Differential expression analysis results for *SOX10*-high versus *SOX10-low* TNBCs from the TGCA and METABRIC expression datasets.

**Table-S4**. Ontology enrichment results for differentially expressed genes from Table-S3.

**Table-S5**. WGCNA module membership data (kTotal, total connectivity across the breast cancer network; kWithin, intra-modular connectivity; kOut, extra-modular connectivity; kDiff, kWithin-kOut; kME, module eigengene correlations; p, p-values).

**Table-S6**. Module eigengene values for TCGA cases.

**Table-S7**. Module eigengene values for METABRIC cases.

**Table-S8**. Module preservation results for the ICGC RNAseq dataset.

**Table-S9**. Module membership ontology enrichment results.

**Table-S10**. SOXE-module network metrics: eigencentrality, betweenness centrality, kWithin, submodule membership and overlap with SOX10’s normal breast module.

**Table-S11**. Neural crest genesets used in this study.

**Table-S12**. WGCNA module membership data (kTotal, total connectivity across the normal breast sample network; kWithin, intra-modular connectivity; kOut, extra-modular connectivity; kDiff, kWithin-kOut; kME, module eigengene correlations; p, p-values)

**Table-S13**. Gene ontology enrichment for genes exclusive to SOX10’s normal module or the SOXE-module, and 109 overlapping genes.

**Table-S14**. Gene ontology enrichment for methylation clusters a-c.

## References

1. Fulford, L.G., et al., Basal-like grade III invasive ductal carcinoma of the breast: patterns of metastasis and long-term survival. Breast Cancer Res, 2007. 9(1): p. R4.

2. Prat, A., et al., Phenotypic and molecular characterization of the claudin-low intrinsic subtype of breast cancer. Breast cancer research: BCR, 2010. 12(5): p. R68.

3. Symmans, W.F., et al., Long-Term Prognostic Risk After Neoadjuvant Chemotherapy Associated With Residual Cancer Burden and Breast Cancer Subtype. J Clin Oncol, 2017. 35(10): p. 1049–1060.

4. Gao, R., et al., Punctuated copy number evolution and clonal stasis in triple-negative breast cancer. Nat Genet, 2016. 48(10): p. 1119–30.

5. Wang, Y., et al., Clonal evolution in breast cancer revealed by single nucleus genome sequencing. Nature, 2014. 512(7513): p. 155–60.

6. Yates, L.R., et al., Subclonal diversification of primary breast cancer revealed by multiregion sequencing. Nat Med, 2015. 21(7): p. 751–9.

7. Yang, F., et al., Intratumor heterogeneity predicts metastasis of triple-negative breast cancer. Carcinogenesis, 2017. 38(9): p. 900–909.

8. Lin, B., et al., Modulating Cell Fate as a Therapeutic Strategy. Cell Stem Cell, 2018. 23(3): p. 329–341.

9. Nguyen, D.X., P.D. Bos, and J. Massagué, Metastasis: from dissemination to organ-specific colonization. Nature Reviews Cancer, 2009. 9(4): p. 274–284.

10. Gupta, P.B., et al., Phenotypic Plasticity: Driver of Cancer Initiation, Progression, and Therapy Resistance. Cell Stem Cell, 2019. 24(1): p. 65–78.

11. Hinohara, K. and K. Polyak, Intratumoral Heterogeneity: More Than Just Mutations. Trends Cell Biol, 2019. 29(7): p. 569–579.

12. Bell, C.C. and O. Gilan, Principles and mechanisms of non-genetic resistance in cancer. Br J Cancer, 2020. 122(4): p. 465–472.

13. Granit, R.Z., et al., Regulation of Cellular Heterogeneity and Rates of Symmetric and Asymmetric Divisions in Triple-Negative Breast Cancer. Cell Rep, 2018. 24(12): p. 3237–3250.

14. Keller, P.J., et al., Defining the cellular precursors to human breast cancer. Proc Natl Acad Sci U S A, 2012. 109(8): p. 2772–7.

15. Lim, E., et al., Aberrant luminal progenitors as the candidate target population for basal tumor development in BRCA1 mutation carriers. Nature Medicine, 2009. 15(8): p. 907–913.

16. Molyneux, G., et al., BRCA1 basal-like breast cancers originate from luminal epithelial progenitors and not from basal stem cells. Cell Stem Cell, 2010. 7(3): p. 403–17.

17. Proia, T.A., et al., Genetic predisposition directs breast cancer phenotype by dictating progenitor cell fate. Cell Stem Cell, 2011. 8(2): p. 149–63.

18. Chaffer, C.L., et al., Normal and neoplastic nonstem cells can spontaneously convert to a stem-like state. Proceedings of the National Academy of Sciences of the United States of America, 2011.108(19): p. 7950–5.

19. Hinohara, K., et al., KDM5 Histone Demethylase Activity Links Cellular Transcriptomic Heterogeneity to Therapeutic Resistance. Cancer Cell, 2018. 34(6): p. 939-953 e9.

20. Risom, T., et al., Differentiation-state plasticity is a targetable resistance mechanism in basal-like breast cancer. Nat Commun, 2018. 9(1): p. 3815.

21. Flavahan, W.A., E. Gaskell, and B.E. Bernstein, Epigenetic plasticity and the hallmarks of cancer. Science, 2017. 357(6348).

22. Stirzaker, C., et al., Methylome sequencing in triple-negative breast cancer reveals distinct methylation clusters with prognostic value. Nat Commun, 2015. 6: p. 5899.

23. Deblois, G., et al., Epigenetic switch-induced viral mimicry evasion in chemotherapy resistant breast cancer. Cancer Discov, 2020.

24. Dravis, C., et al., Epigenetic and Transcriptomic Profiling of Mammary Gland Development and Tumor Models Disclose Regulators of Cell State Plasticity. Cancer Cell, 2018. 34(3): p. 466-482 e6.

25. Hu, N., P.H. Strobl-Mazzulla, and M.E. Bronner, Epigenetic regulation in neural crest development. Dev Biol, 2014. 396(2): p. 159–68.

26. Southard-Smith, E.M., L. Kos, and W.J. Pavan, Sox10 mutation disrupts neural crest development in Dom Hirschsprung mouse model. Nat Genet, 1998. 18(1): p. 60–4.

27. Kim, J., et al., SOX10 maintains multipotency and inhibits neuronal differentiation of neural crest stem cells. Neuron, 2003. 38(1): p. 17–31.

28. McKeown, S.J., et al., Sox10 overexpression induces neural crest-like cells from all dorsoventral levels of the neural tube but inhibits differentiation. Dev Dyn, 2005. 233(2): p. 430–44.

29. Dravis, C., et al., Sox10 Regulates Stem/Progenitor and Mesenchymal Cell States in Mammary Epithelial Cells. Cell Rep, 2015. 12(12): p. 2035–48.

30. Chen, Z., et al., FGF signaling activates a Sox9-Sox10 pathway for the formation and branching morphogenesis of mouse ocular glands. Development, 2014. 141(13): p. 2691–701.

31. Athwal, H.K., et al., Sox10 Regulates Plasticity of Epithelial Progenitors toward Secretory Units of Exocrine Glands. Stem Cell Reports, 2019. 12(2): p. 366–380.

32. Guo, W., et al., Slug and Sox9 cooperatively determine the mammary stem cell state. Cell, 2012. 148(5): p. 1015–28.

33. Mertelmeyer, S., et al., The transcription factor Sox10 is an essential determinant of branching morphogenesis and involution in the mouse mammary gland. Sci Rep, 2020. 10(1): p. 17807.

34. Kim, Y.J., et al., Generation of multipotent induced neural crest by direct reprogramming of human postnatal fibroblasts with a single transcription factor. Cell Stem Cell, 2014. 15(4): p. 497–506.

35. Ivanov, S.V., et al., Diagnostic SOX10 gene signatures in salivary adenoid cystic and breast basal-like carcinomas. Br J Cancer, 2013. 109(2): p. 444–51.

36. Panaccione, A., et al., Expression Profiling of Clinical Specimens Supports the Existence of Neural Progenitor-Like Stem Cells in Basal Breast Cancers. Clin Breast Cancer, 2017. 17(4): p. 298-306 e7.

37. Cimino-Mathews, A., et al., Neural crest transcription factor Sox10 is preferentially expressed in triple-negative and metaplastic breast carcinomas. Hum Pathol, 2013. 44(6): p. 959–65.

38. Jamidi, S.K., et al., SOX10 as a sensitive marker for triple negative breast cancer. Histopathology, 2020.

39. Burstein, M.D., et al., Comprehensive genomic analysis identifies novel subtypes and targets of triple-negative breast cancer. Clin Cancer Res, 2015. 21(7): p. 1688–98.

40. Hu, N., et al., DNA methyltransferase 3B regulates duration of neural crest production via repression of Sox10. Proc Natl Acad Sci U S A, 2014. 111(50): p. 17911–6.

41. Strobl-Mazzulla, P.H. and M.E. Bronner, A PHD12-Snail2 repressive complex epigenetically mediates neural crest epithelial-to-mesenchymal transition. J Cell Biol, 2012. 198(6): p. 999–1010.

42. Pellacani, D., et al., Analysis of Normal Human Mammary Epigenomes Reveals Cell-Specific Active Enhancer States and Associated Transcription Factor Networks. Cell Rep, 2016. 17(8): p. 2060–2074.

43. Curtis, C., et al., The genomic and transcriptomic architecture of 2,000 breast tumours reveals novel subgroups. Nature, 2012.

44. TCGA, Cancer Genome Atlas Network: Comprehensive molecular portraits of human breast tumours. Nature, 2012. 490(7418): p. 61–70.

45. Nik-Zainal, S., et al., Landscape of somatic mutations in 560 breast cancer whole-genome sequences. Nature, 2016.

46. Daemen, A., et al., Modeling precision treatment of breast cancer. Genome Biol, 2013. 14(10): p. R110.

47. Neve, R.M., et al., A collection of breast cancer cell lines for the study of functionally distinct cancer subtypes. Cancer Cell, 2006. 10(6): p. 515–527.

48. McCart Reed, A.E., et al., The Brisbane Breast Bank. Open Journal of Bioresources, 2018. 5: p. 5.

49. Saunus, J.M., et al., Multidimensional phenotyping of breast cancer cell lines to guide preclinical research. Breast Cancer Res Treat, 2018.

50. Qi, J., et al., SOX10 – A Novel Marker for the Differential Diagnosis of Breast Metaplastic Squamous Cell Carcinoma. Cancer Manag Res, 2020. 12: p. 4039–4044.

51. McCart Reed, A.E., et al., Phenotypic and molecular dissection of metaplastic breast cancer and the prognostic implications. J Pathol, 2019. 247(2): p. 214–227.

52. Zhang, B. and S. Horvath, A general framework for weighted gene co-expression network analysis. Stat Appl Genet Mol Biol, 2005. 4: p. Article17.

53. Langfelder, P. and S. Horvath, WGCNA: an R package for weighted correlation network analysis. BMC Bioinformatics, 2008. 9: p. 559.

54. Denkert, C., et al., Tumour-infiltrating lymphocytes and prognosis in different subtypes of breast cancer: a pooled analysis of 3771 patients treated with neoadjuvant therapy. Lancet Oncol, 2018. 19(1): p. 40–50.

55. Simoes-Costa, M., et al., Transcriptome analysis reveals novel players in the cranial neural crest gene regulatory network. Genome Res, 2014. 24(2): p. 281–90.

56. Pellacani, D., et al., Transcriptional regulation of normal human mammary cell heterogeneity and its perturbation in breast cancer. EMBO J, 2019. 38(14): p. e100330.

57. Davies, H., et al., HRDetect is a predictor of BRCA1 and BRCA2 deficiency based on mutational signatures. Nat Med, 2017. 23(4): p. 517–525.

58. Hayano, M., et al., DNA Break-Induced Epigenetic Drift as a Cause of Mammalian Aging. bioRxiv, 2019: p. 808659.

59. Yang, J.-H., et al., Erosion of the Epigenetic Landscape and Loss of Cellular Identity as a Cause of Aging in Mammals. bioRxiv, 2019: p. 808642.

60. Zhou, W., et al., DNA methylation loss in late-replicating domains is linked to mitotic cell division. Nat Genet, 2018. 50(4): p. 591–602.

61. Medvedeva, Y.A., et al., EpiFactors: a comprehensive database of human epigenetic factors and complexes. Database (Oxford), 2015. 2015: p. bav067.

62. Jin, C., et al., TET1 is a maintenance DNA demethylase that prevents methylation spreading in differentiated cells. Nucleic Acids Res, 2014. 42(11): p. 6956–71.

63. Putiri, E.L., et al., Distinct and overlapping control of 5-methylcytosine and 5-hydroxymethylcytosine by the TET proteins in human cancer cells. Genome Biol, 2014. 15(6): p. R81.

64. Good, C.R., et al., TET1-Mediated Hypomethylation Activates Oncogenic Signaling in Triple-Negative Breast Cancer. Cancer Res, 2018. 78(15): p. 4126–4137.

65. Timp, W., et al., Large hypomethylated blocks as a universal defining epigenetic alteration in human solid tumors. Genome Med, 2014. 6(8): p. 61.

66. Hovestadt, V., et al., Decoding the regulatory landscape of medulloblastoma using DNA methylation sequencing. Nature, 2014. 510(7506): p. 537–41.

67. Foroutan, M., et al., Single sample scoring of molecular phenotypes. BMC Bioinformatics, 2018. 19(1): p. 404.

68. Drier, Y., M. Sheffer, and E. Domany, Pathway-based personalized analysis of cancer. Proc Natl Acad Sci U S A, 2013. 110(16): p. 6388–93.

69. Livshits, A., et al., Pathway-based personalized analysis of breast cancer expression data. Mol Oncol, 2015. 9(7): p. 1471–83.

70. Edgar, R., et al., LifeMap Discovery: the embryonic development, stem cells, and regenerative medicine research portal. PLoS One, 2013. 8(7): p. e66629.

71. Nonaka, D., L. Chiriboga, and B.P. Rubin, Sox10: a pan-schwannian and melanocytic marker. Am J Surg Pathol, 2008. 32(9): p. 1291–8.

72. Feinberg, A.P., R. Ohlsson, and S. Henikoff, The epigenetic progenitor origin of human cancer. Nat Rev Genet, 2006. 7(1): p. 21–33.

73. Wahl, G.M. and B.T. Spike, Cell state plasticity, stem cells, EMT, and the generation of intra-tumoral heterogeneity. NPJ Breast Cancer, 2017. 3: p. 14.

74. Visvader, J.E. and J. Stingl, Mammary stem cells and the differentiation hierarchy: current status and perspectives. Genes Dev, 2014. 28(11): p. 1143–58.

75. Liao, C. and Q. Zhang, BBOX1 promotes triple-negative breast cancer progression by controlling IP3R3 stability. Mol Cell Oncol, 2020. 7(6): p. 1813526.

76. Liao, C., et al., Identification of BBOX1 as a Therapeutic Target in Triple-Negative Breast Cancer. Cancer Discov, 2020. 10(11): p. 1706–1721.

77. Zhu, L., et al., BCL11A enhances stemness and promotes progression by activating Wnt/beta-catenin signaling in breast cancer. Cancer Manag Res, 2019. 11: p. 2997–3007.

78. Errico, A., Genetics: BCL11A-targeting triple-negative breast cancer? Nat Rev Clin Oncol, 2015. 12(3): p. 127.

79. Khaled, W.T., et al., BCL11A is a triple-negative breast cancer gene with critical functions in stem and progenitor cells. Nat Commun, 2015. 6: p. 5987.

80. Saggese, P., et al., Metabolic Regulation of Epigenetic Modifications and Cell Differentiation in Cancer. Cancers (Basel), 2020. 12(12).

81. Miettinen, M., et al., Sox10--a marker for not only schwannian and melanocytic neoplasms but also myoepithelial cell tumors of soft tissue: a systematic analysis of 5134 tumors. Am J Surg Pathol, 2015. 39(6): p. 826–35.

82. A, A.A., et al., TFAP2B, AP-1 and JAZF1 Expression in Tissues of Papillary Thyroid Carcinoma Patients; Clinical, Pathological and Prognostic Values. Asian Pac J Cancer Prev, 2020. 21(8): p. 2415–2421.

83. Huang, J. and L. Guo, Knockdown of SOX9 Inhibits the Proliferation, Invasion, and EMT in Thyroid Cancer Cells. Oncol Res, 2017. 25(2): p. 167–176.

84. Schulten, H.J., et al., Comparison of microarray expression profiles between follicular variant of papillary thyroid carcinomas and follicular adenomas of the thyroid. BMC Genomics, 2015. 16 Suppl 1: p. S7.

85. Nakayama, T., et al., Expression of the ets-1 proto-oncogene in human thyroid tumor. Mod Pathol, 1999. 12(1): p. 61–8.

86. Bronchain, O.J., et al., Implication of thyroid hormone signaling in neural crest cells migration: Evidence from thyroid hormone receptor beta knockdown and NH3 antagonist studies. Mol Cell Endocrinol, 2017. 439: p. 233–246.

87. Simoes-Costa, M. and M.E. Bronner, Establishing neural crest identity: a gene regulatory recipe. Development, 2015. 142(2): p. 242–57.

88. Triolo, V.A., Nineteenth Century Foundations of Cancer Research Advances in Tumor Pathology, Nomenclature, and Theories of Oncogenesis. Cancer Res, 1965. 25: p. 75–106.

89. Waddington, C.H., Cancer and the Theory of Organisers. Nature, 1935. 135(3416): p. 606–608.

90. Kulesa, P.M., et al., Reprogramming metastatic melanoma cells to assume a neural crest cell-like phenotype in an embryonic microenvironment. Proc Natl Acad Sci U S A, 2006. 103(10): p. 3752–7.

91. Ben-Porath, I., et al., An embryonic stem cell-like gene expression signature in poorly differentiated aggressive human tumors. Nat Genet, 2008. 40(5): p. 499–507.

