## Supplementary material for "Epigenome erosion drives neural crest-like phenotypic mimicry in triple-negative breast cancer and other SOX10+ malignancies": Fig-S

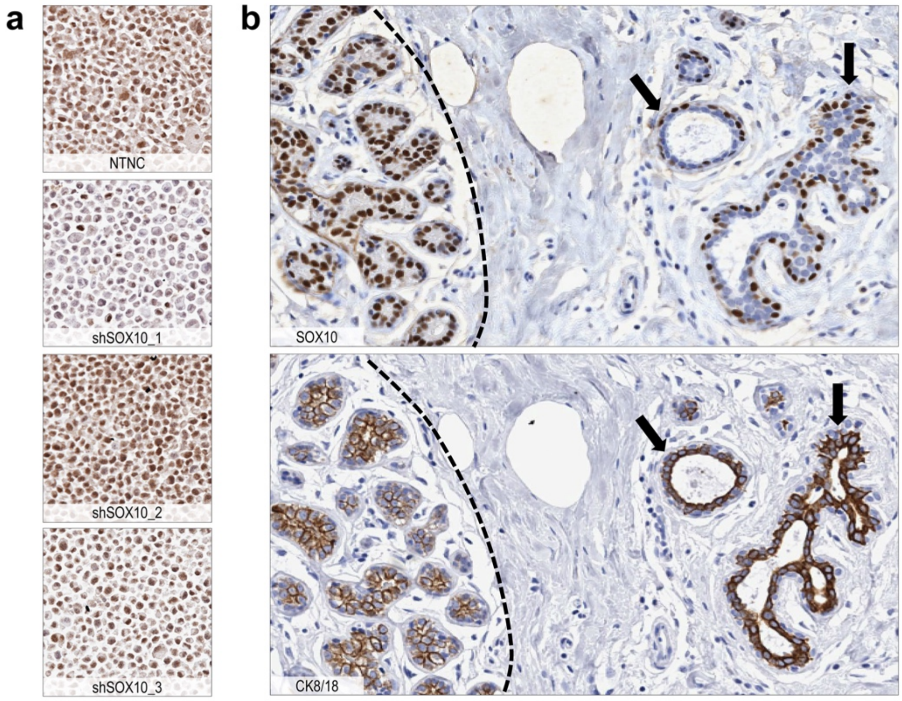


**Figure-S1: Data supporting Fig-1.**

(**a**) Validation of SOX10 antibody specificity in FFPE MDA-MB-435 cell pellet sections. Prior to fixation and analysis, cells were stably transduced with non-template negative control (NTNC) or *SOX10* shRNAs. (**b**) IHC analysis of SOX10 and cytokeratin (CK) 8/18 expression in serial RM tissue sections. Representative image shows a lobule with SOX10+ luminal epithelia and weak expression of CK8/18 (outlined); and nearby ducts (arrows) in which the luminal epithelium lacks SOX10 expression but stains strongly for CK8/18.

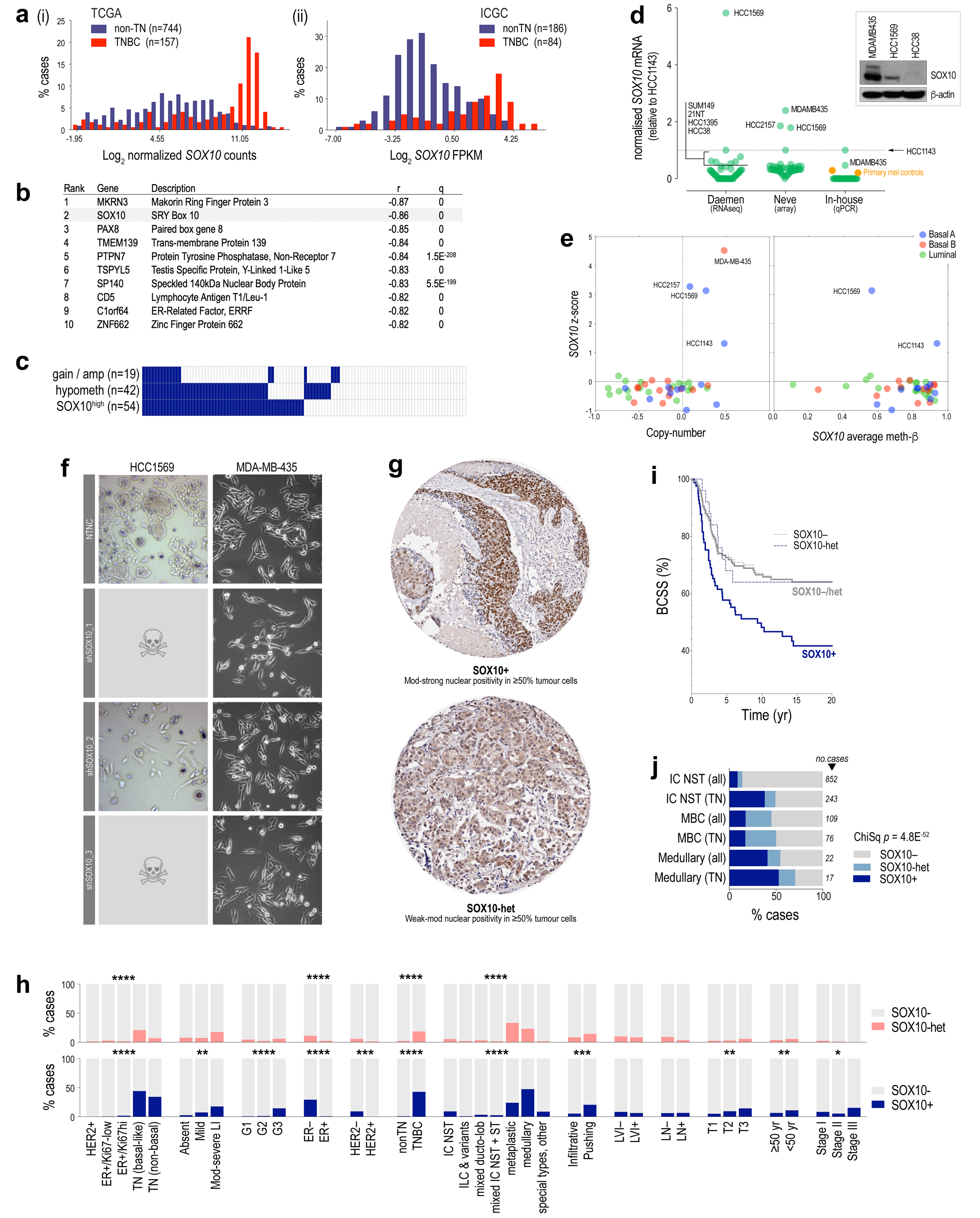


**Figure-S2: Data supporting Fig-2.**

(**a**) *SOX10* mRNA distribution in TNBC and non-TN cases from TCGA and ICGC cohorts. (**b**) Of all genes expressed in breast cancer, *SOX10* has the second strongest relationship with gene methylation (Broad Institute TCGA Genome Data Analysis Centre). (**c**) Percentages of *SOX10* hypomethylated (90$^{th}$ percentile of melanoma values) and copy-number altered cases (GISTIC). *amp, amplified*. (**d**) Expression of *SOX10* mRNA in breast cancer cell lines; three independent datasets shown. Inset: protein expression in selected lines determined by Western analysis (antibody validation in Fig-S1). (**e**) Relationships between *SOX10* expression, copy-number and gene-averaged methylation beta values in breast cancer cell lines. The high levels of promoter methylation and barely detectable mRNA in the majority of lines suggested that *SOX10* silencing is a cause and/or consequence of *in vitro* adherent selection. Culturing a range of lines as non-adherent tumourspheres was not sufficient to reactivate SOX10 expression (data not shown). (**f**) Light microscopy images (20x magnification) comparing cell morphology and cell death (☠) of MDA-MB-435 melanoma and HCC1569 basal-like breast cancer cell lines stably transduced with *SOX10*-targeted shRNA, or a non-targeted negative control (NTNC) hairpin. HCC1569-sh*SOX10* derivatives did not survive beyond three passages after antibiotic selection. MDA-MB-435 were viable over multiple (>10) passages but assumed more mesenchymal morphology. (**g**) Additional IHC images showing strong positivity, heterogeneous staining and tumour-associated normal staining in TNBC samples. The heterogeneous core also exemplifies cytoplasmic staining, which was detected in both ER+ and ER– cases and was not associated with any of the clinico-pathologic variables (Table-S2) that we analysed (data not shown). (**h**) Proportions of SOX10-neg versus SOX10-heterogeneous (upper panel) and positive (lower panel). Chi Square test p-values are shown: *$p<0.01$; **$p<0.001$; ***$p<0.0001$; ****$p<0.00001$. (**i**) Overlapping survival curves for SOX10-neg and heterogeneous staining in TNBC. (**j**) SOX10 heterogeneity in metaplastic and medullary TNBCs compared to tumours of no special type (NST).

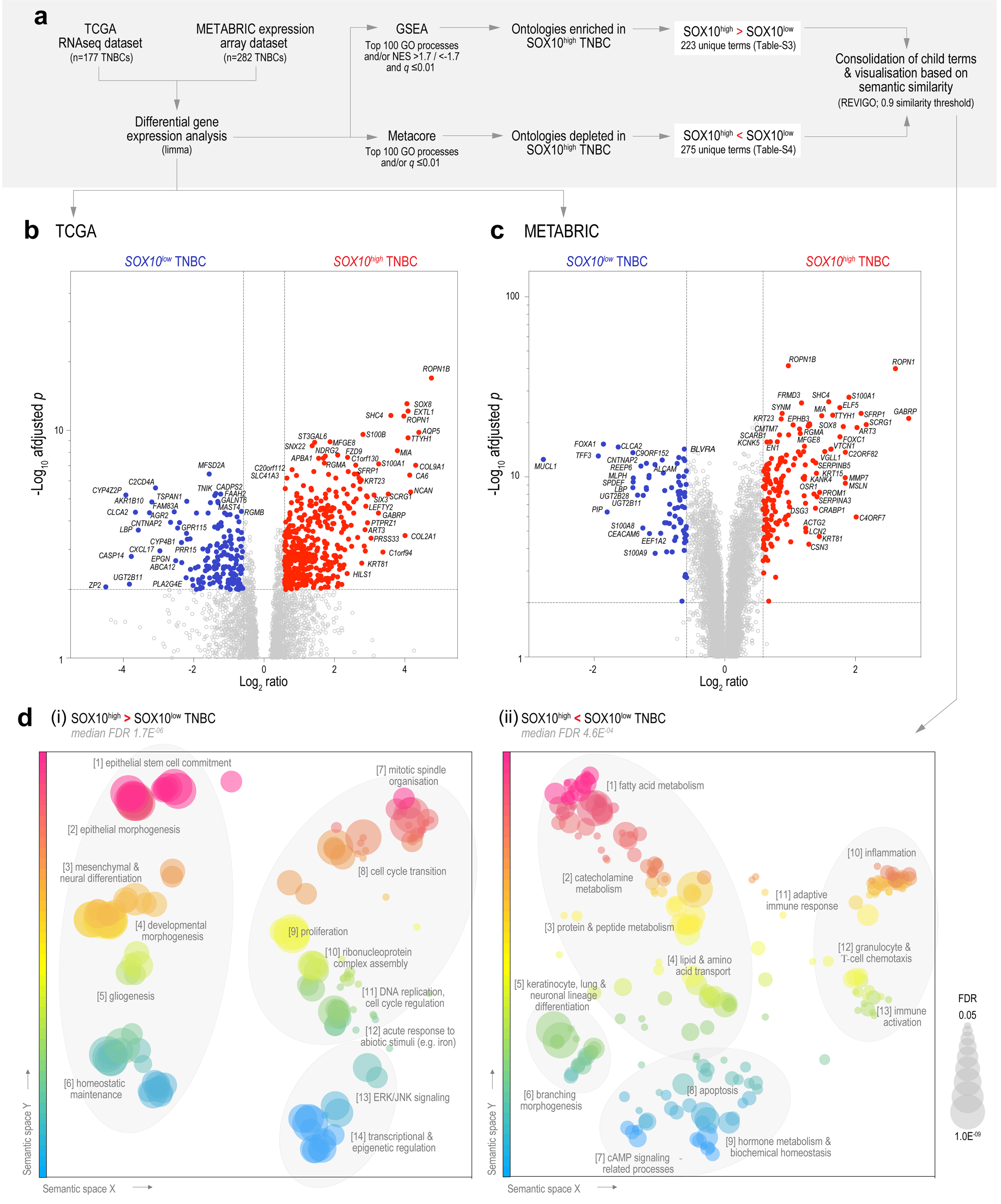


**Figure-S3: Gene ontology profiles of SOX10-high and –low TNBCs.**

(**a**) TNBC expression analysis strategy. (**b/c**) Volcano plots indicate transcripts with high significance in differential expression analysis (also see Table-S3). (**d**) Semantic similarity plots summarising major gene ontologies (GO) enriched in *SOX10*-high and -low TNBCs. Differential gene expression analysis was performed separately on TCGA and METABRIC TNBC datasets. Up to 100 of the most significant (corrected *p*≤0.05) GO processes identified through enrichment analysis (MetaCore and GSEA) were clustered in 2D semantic space using REVIGO (‘reduce & visualise gene ontology’), which removes redundancy and assigns X-Y coordinates on the basis of semantic similarity analysis of pre-computed information content[1]. GO terms are coloured according to Y coordinates and sized proportionally to the corrected enrichment p-value (FDR, false discovery rate).

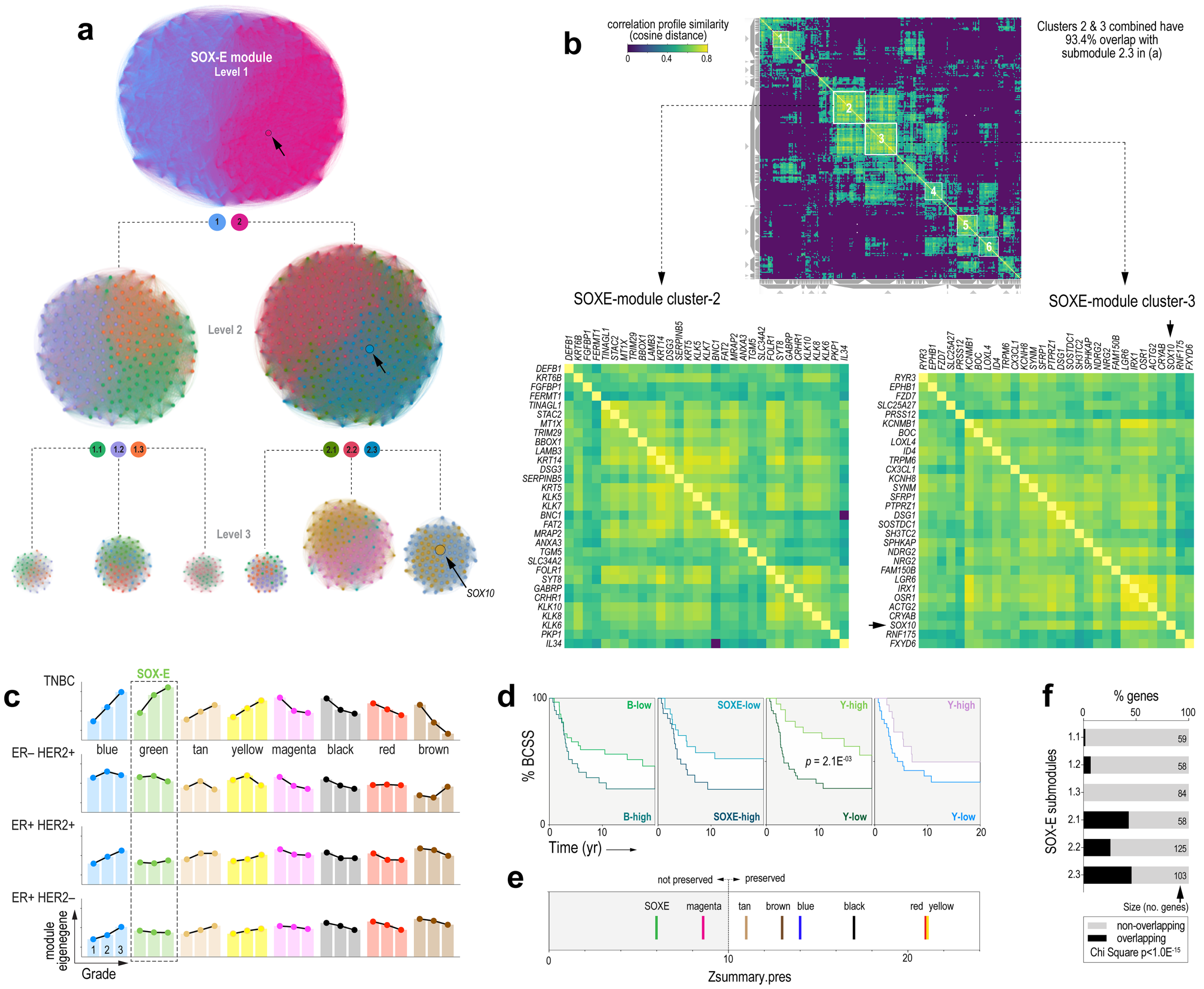


**Figure-S4: Data supporting Fig-3.** (**a/b**) Identification of SOXE-module hubs using community detection algorithms to analyse topological structure (a) and unsupervised clustering to map gene correlation profile similarity (according to cosine distance) (TCGA datasets). According to these complementary approaches, approximately 60 genes are most essential to SOXE-module architecture and information flow (89% of submodule 2.3 genes from (a) cluster together in (b). Conversely, 93.4% of clusters 1+2 genes from (b) are located within submodule 2.3 from (a). See also Table-S12. (**c**) Relationship between SOXE-module expression and histological grade in breast cancer (METABRIC dataset). (**d**) Kaplan Meier analysis of the effects of module co-expression in TNBCs from patients treated with chemotherapy and/or radiotherapy (METABRIC dataset; treatment subpopulation from the cohort in Fig-3g). BCSS, breast cancer-specific survival. ME fraction thresholds for classifying cases as high or low were 0.33 for SOXE/blue and 0.1 for yellow. (**e**) Breast cancer module preservation in normal breast samples (n=50, TCGA). The z-score threshold considered to represent preservation is 10 (indicated). (**f**) Proportions of SOXE-submodule genes shared with *SOX10's* normal breast module, showing significant enrichment in submodules 2.1 and 2.3 in particular

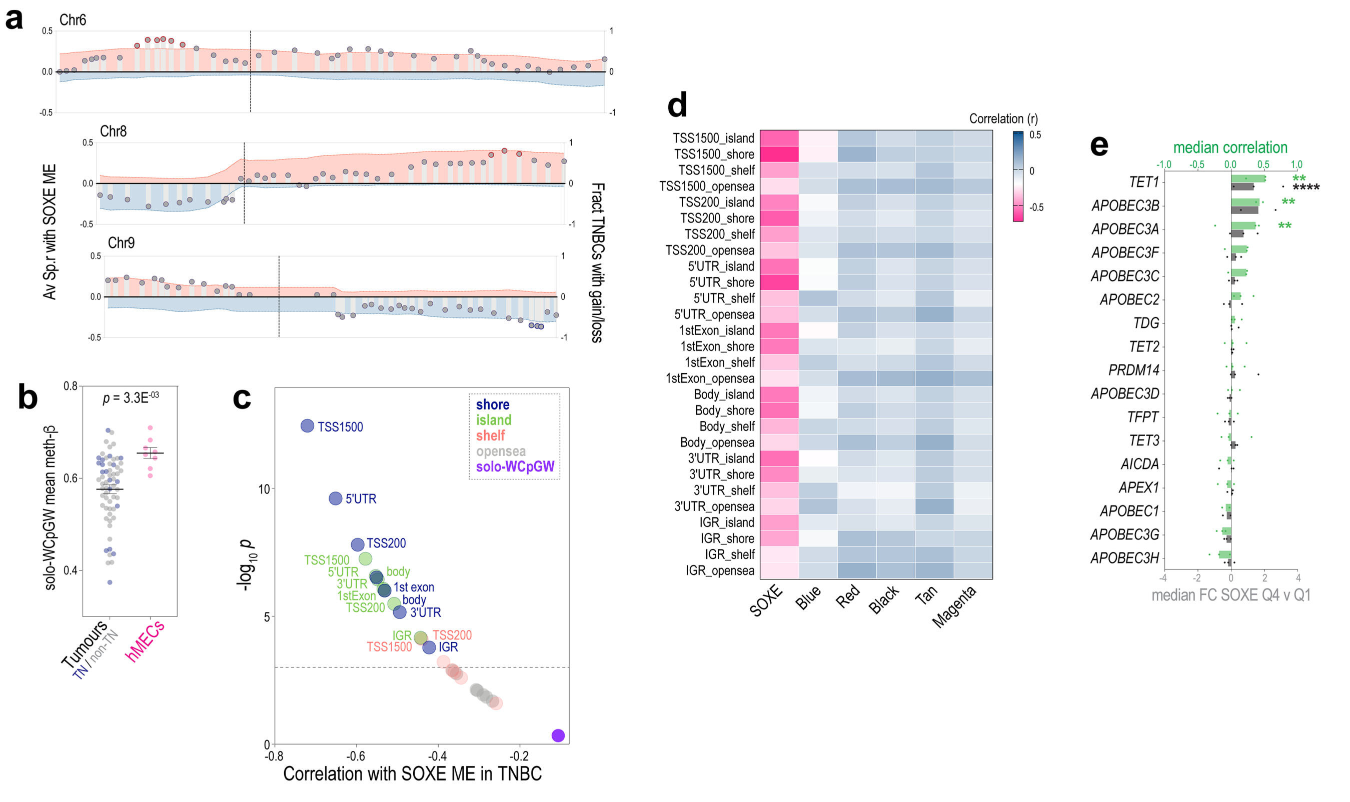


**Figure-S5: Data supporting Fig-5.**

(**a**) Relationship between SOXE-ME values and large-scale copy-number alterations (CNAs) on chromosomes 6, 8 and 9 (TCGA datasets). X-axis, chromosome position, to scale; left y-axis and bars: spearman correlation between SOXE-ME values and genes, averaged across cytobands; right y-axis and shaded area: average copy-number gain/amplification (red) and loss/deletion (blue) for genes averaged across cytobands; dotted lines, centromere position. Red/blue circles highlight loci with the highest and lowest correlation coefficients, which generally coincide with TNBC's most frequently gained/lost regions, respectively. (**b**) Average Illumina EPIC 850k methylation array beta values for solo-WCpGW sites in breast tumours versus FACS-sorted hMEC subtypes (see ref[2] and Fig-1). Stats: Mann-Whitney test. **(c)** Correlation (x-axis) and significance (y) of methylation beta values (averaged for the categories listed) versus SOXE-ME values in TNBC. **(d)** Complete correlation matrix for all methylation categories and all WGCNA modules expressed in TNBC (TCGA dataset). (**e**) Relationships between expression of the SOXE module and demethylases in the EpiFactors database[3]. Plots summarise average Spearman correlations between the expression of demethylating enzymes and SOXE ME values (y-axis), and average fold-change (FC; Mann-Whitney test) between TNBCs expressing high (quartile-4) and low (quartile-1) levels of the demethylases (n=3 expression datasets: TCGA, METABRIC, ICGC). **$p$<0.01; ****$p$<0.0001.

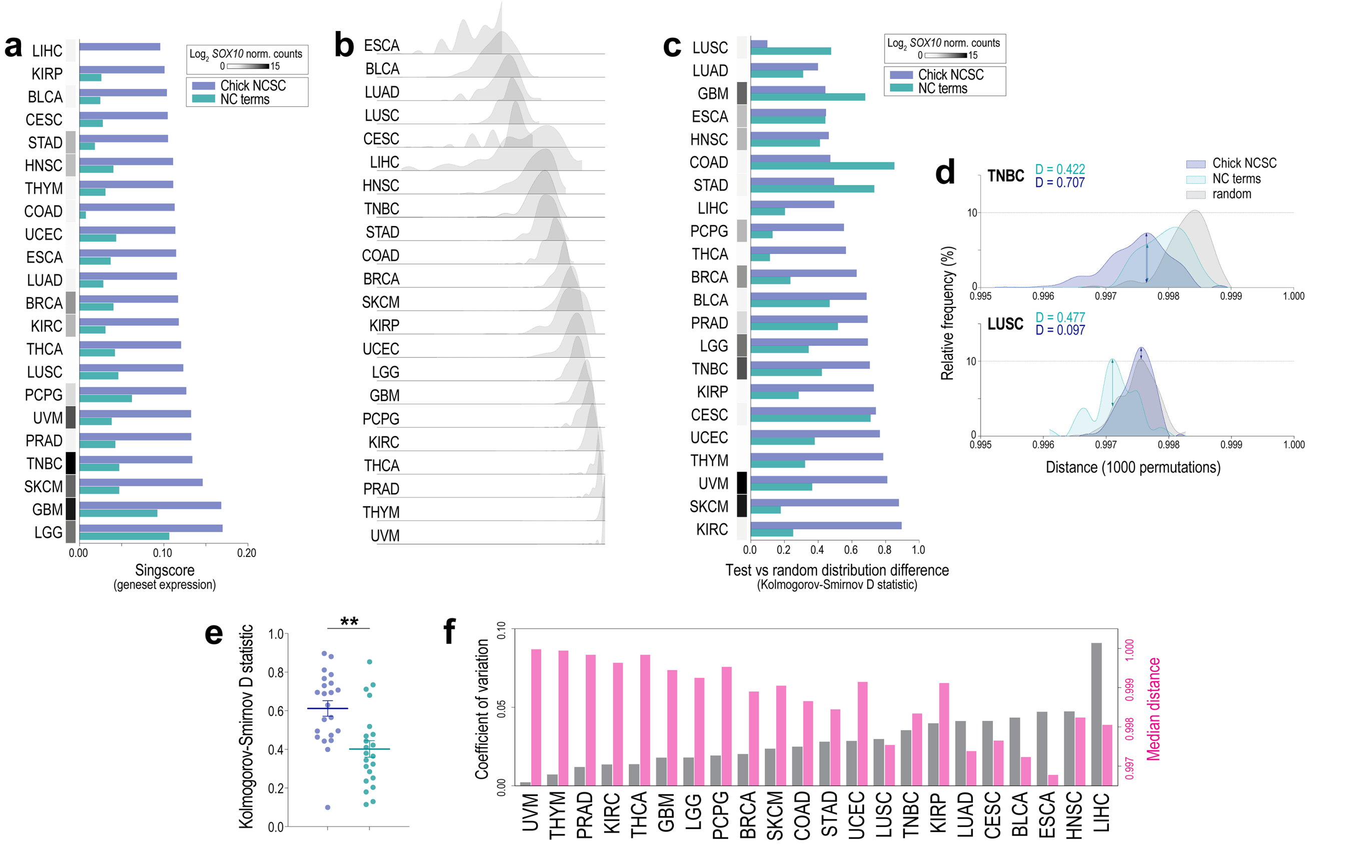


**Figure-S6: Data supporting Fig-7.**

(**a**) NCSC geneset expression and *SOX10* normalised RNAseq counts in 22 different malignancies with RNAseq dataset available from TCGA according to *singscore*. (**b**) Representation of the variability amongst cancer-specific transcriptome network structures. Pairwise correlation matrices were built from RNAseq data for each malignancy, then sampled 1000 times (1 sample = median pairwise distance amongst 60 gene pairs selected at random), and the distributions of these measurements were plotted, with splines fitted universally for visualisation. The y-axis is frequency, while the x-axis is median inter-gene distance, with lower values indicating closer proximity within the network. Narrow distributions (e.g., UVM, THYM) represent more homogeneous network architecture, while broad and/or uneven distributions (e.g., ESCA, BLCA) represent heterogeneous network architecture. (**c**) Median inter-gene distances for two independent NCSC genesets were consistently lower than the control (median pairwise distance amongst 60 gene pairs selected either from the NCSC genesets or at random as a control; 1000 permutations for each of the three genesets in each malignancy). The plot compares *SOX10* expression with the extent of these differences, represented by the Kolmogorov-Smirnov D statistic (see (**d**)). (**e**) Comparison of KS-D values for NC-terms and ch.NCSC genesets. Stats: Mann-Whitney test ($p$<0.01). (**f**) Median distance versus co-efficient of variation for random control permutations, highlighting the different degrees of variability across the cancer-specific networks (complements (**b**)).

**Abbreviations**: BLCA, Bladder Urothelial Carcinoma; BRCA, Breast invasive carcinoma (non-TN cases); CESC, Cervical squamous cell carcinoma and endocervical adenocarcinoma; COAD, Colon adenocarcinoma; ESCA, Oesophageal carcinoma; GBM, Glioblastoma multiforme; HNSC, Head and Neck squamous cell carcinoma; KIRC, Kidney renal clear cell carcinoma; KIRP, Kidney renal papillary cell carcinoma; LGG, Brain Lower Grade Glioma; LIHC, Liver hepatocellular carcinoma; LUAD, Lung adenocarcinoma; LUSC, Lung squamous cell carcinoma; PCPG, Pheochromocytoma and Paraganglioma; PRAD, Prostate adenocarcinoma; SKCM, Skin Cutaneous Melanoma; STAD, Stomach adenocarcinoma; THCA, Thyroid carcinoma; THYM, Thymoma; UCEC, Uterine Corpus Endometrial Carcinoma; UVM, Uveal Melanoma.

### References


1. Supek, F., et al., *REVIGO summarizes and visualizes long lists of gene ontology terms.* PLoS One, 2011. **6**(7): p. e21800.

2. Nones, K., et al., *Whole-genome sequencing reveals clinically relevant insights into the aetiology of familial breast cancers.* Ann Oncol, 2019. **30**(7): p. 1071-1079.

3. Medvedeva, Y.A., et al., *EpiFactors: a comprehensive database of human epigenetic factors and complexes.* Database (Oxford), 2015. **2015**: p. bav067.
