## supplementary file 1 for "Epigenome erosion drives neural crest-like phenotypic mimicry in triple-negative breast cancer and other SOX10+ malignancies"

### MATERIALS, METHODS & DATA AVAILABILITY

##### *Human tissue samples*

Reduction mammoplasty (RM) samples were obtained in collaboration with Dr William Cockburn (Wesley Hospital, Brisbane) and the Royal Brisbane and Women's Hospital (RBWH) Plastics Unit. Nineteen RM specimens were used for IHC and IF analysis, and two for methylation arrays. Age, parity and menopausal status of these patients were unknown. 30% of cases showed fibrocystic change and 10% presented with columnar cell lesions (histopathology review by SRL). The following breast tumour cohorts were also used:

1. The Queensland follow-up (QFU) cohort[1,2]: clinically annotated, archival breast tumour samples from patients treated by the RBWH Breast Unit between 1987 and 1994. Tumours were sampled as duplicate cores in tissue microarrays (TMAs).
2. Nottingham University Hospital early-stage breast cancer cohort[3,4]: archival breast tumour samples with clinicopathologic annotation including at least 15 years' follow-up (TMA format).
3. Tumour-associated normal breast tissue from 32 QFU cohort cases (whole sections).
4. Primary TNBCs with patient-matched brain metastases (TMA format).
5. FFPE metaplastic carcinomas from the Asia Pacific Metaplastic Breast Cancer Consortium[5] (n=105; whole sections.)

##### *Immunohistochemistry (IHC)*

Formalin-fixed, paraffin-embedded (FFPE) tissue samples or TMAs were sectioned, deparaffinised, subjected to antigen retrieval and chromogenically stained as described[6] and detailed in Table-M1. Slides were scanned using the Aperio ScanScope T2 digital scanning system at 40x magnification for digital scoring. TMA images were segmented using Spectrum software (Aperio), and high-resolution images of individual cores were extracted and scored by two observers in a blinded fashion (hidden metadata tags corresponding to TMA position were used to link clinical and sample data).

##### *Immunofluorescence (IF)*

FFPE RM tissue sections were sectioned, deparaffinised, subjected to antigen retrieval and stained as described[7] (Table-M1). Briefly, primary antibodies diluted in tris-buffered saline (TBS) were incubated on tissue sections for 1 h at room temperature, washed in TBS then incubated with secondary antibodies for 30 min in the dark. To minimise tissue autofluorescence, slides were stained with SUDAN Black for 20 min in the dark (Sigma #S-2380), then washed (0.1% TBS-Tween (30 min), TBS (10 min). Slides were mounted using Vectashield (Vecta Labs) with DAPI (Sigma-Aldrich), cover-slipped, sealed and imaged on a Carl Zeiss MicroImaging system using Axio Vision LE version 4.8.2 (PerkinElmer).

##### *Fresh reduction mammoplasty (RM) tissue processing and fluorescence-activated cell sorting (FACS)*

RM samples were processed, and single cell suspensions prepared as previously described[7,8]. Briefly, tissue was cut into small pieces (~5 $mm^3$) and digested overnight with agitation at 37°C in DMEM-F12 (Gibco), foetal bovine serum ((FBS), 5%, Gibco), antibiotic/antimycotic (Gibco), Amphotericin B (2.5 µg/mL, Gibco), collagenase type I-A (200 U/mL, Sigma-Aldrich) and Hyaluronidase I-S (100 U/mL, Sigma-Aldrich). Epithelial organoids were obtained by centrifugation (80×g, 1 min), then dissociated to single cell suspensions for 5-10 min in TrypLE (Gibco), followed by Dispase (5 mg/mL, Gibco) and DNAse-I (100 ug/mL, Invitrogen). Enzymatic activity was quenched in ice-cold Hank's Balanced Salt Solution ((HBSS), Gibco) with 2% FBS and cells were filtered through a 40 µm cell strainer (BD Falcon).

Cell concentration and viability were determined using a Countess® automated counter (Invitrogen) with trypan blue and adjusted to 2.0E$^{6}$/mL. Single cell suspensions (typically 30-60 mL) were labelled for 10 min on ice with Sytox™ blue (Invitrogen) plus a cocktail of fluorescent antibody conjugates to discriminate hMEC subsets (negatively gated, non-epithelial 'lineage' markers: CD31, CD45, CD140b; positively gated hMEC markers: CD49f, EpCAM – see Table-M1). Samples were washed (80×g, 2 min), then resuspended in cold HBSS + 2% FBS for FACS. For robust fluorescence compensation and gating of specific hMEC populations, we also tested in parallel small samples stained with isotype control antibodies, and 'fluorescence minus one' negative controls (five samples from which one of the main conjugates was omitted). Fluorescence data acquisition, gate placement and sorting were performed on a BD FACS Aria II instrument (QIMR Berghofer) and sorted cells were collected on ice before being pelleted (80×g, 2 min) and snap-frozen at -70°C.

**Table-M1: Antibodies and immuno-staining conditions used in this study.**

| Antigen | Application | Conjugate | Clone | Isotype | Cat no. | Host | Supplier | Working dilution | Antigen retrieval | Staining localisation scored | Positivity threshold or parameters scored |
|---|---|---|---|---|---|---|---|---|---|---|---|
| ER | IHC, IF | – | 6F11 | | NCL-L-ER-6F11 | M | Novocastra | 100 | C | nuc | Strong+ >1% cells |
| PR | IHC | – | 1A6 | | NCL-L-PGR-312 | M | Novocastra | 200 | C | nuc | Strong+ >1% cells |
| Ki67 | IHC | – | MIB-1 | | M7240 | M | Dako | 200 | C | nuc | Strong+ ≥20% cells |
| CK5/6 | IHC | – | D5/16B4 | | MAB-1620 | M | Chemicon | 400 | C | memb, cyto | Strong+ >1% cells |
| CK14 | IHC | – | LL002 | | NCL-LL002 | M | Novocastra | 40 | C | memb, cyto | Strong+ >1% cells |
| CK8/18 | IHC, IF | – | 5D3 | | NCL-L-CE | M | Novocastra | 100 | C | memb, cyto | Strong+ >1% cells |
| EGFR | IHC | – | 31G7 | | 280005 | M | Invitrogen | 100 | E | memb | Strong+ >1% cells |
| Ki-67 | IHC, IF | – | MIB-1 | | M7240 | M | Dako | 200 | C | nuc | > 20%+ |
| cKIT | IHC, IF | – | polyclonal | | A4502 | R | Dako | 800 | E | memb | Any+ |
| SOX10 | IHC, IF<br>WB | – | sc-17342 | | N-20 | G | Santa Cruz | 100<br>500 | C | nuc, cyto | Any+ |
| ID4 | IHC | – | 82-12 | | BCH-9 | R | Biocheck | 400 | C | nuc | Any+ |
| ELF5 | IHC | – | polyclonal | | sc-376737 | G | Santa Cruz | 200 | E | nuc, cyto | % and intensity (1-3+) |
| PrPC | IHC | – | SAF32 | | 189720 | M | Cayman Chemicals | 50 | none | memb, cyto | Any+ |
| ITGB4 | IHC | – | D8P6C | | 14803 | R | Cell Signaling | 200 | E | memb | % and intensity (1-3+) |
| SOX9 | IHC | – | polyclonal | | AB5535 | R | Merck Millipore | 5000 | C | nuc | Any+ |
| ROPN1B | IHC | – | polyclonal | | HPA052530 | R | Sigma-Aldrich | 200 | E | cyto | Any+ |
| beta actin | WB | – | 8H10D10 | | 3700 | M | Cell Signaling | 5000 | – | – | – |
| G IgG | IF | AF®488 | polyclonal | | A11008 | R | Invitrogen | 500 | – | – | – |
| M IgG1 | IF | AF®594 | polyclonal | | A21125 | G | Invitrogen | 400 | – | – | – |
| CD49f | FACS | PE-Cy™5 | GoH3 | IgG2a κ | 551129 | R | BD | 100 | – | memb | thresholded in each expt |
| CD140b | FACS | PE | 28D4 | IgG2a κ | 558821 | M | BD | 100 | – | memb | thresholded in each expt |
| CD45 | FACS | PE | HI30 | IgG1 κ | 555483 | M | BD | 100 | – | memb | thresholded in each expt |
| – | FACS | FITC | X40 | IgG1 κ | 349041 | M | BD | matched to test Ab for each lot | | | thresholded in each expt |
| – | FACS | PE-Cy™5 | R35-95 | IgG2a κ | 551066 | R | BD | matched to test Ab for each lot | | | thresholded in each expt |
| – | FACS | PE | X39 | IgG2a κ | 349053 | M | BD | matched to test Ab for each lot | | | thresholded in each expt |
| – | FACS | PE | MOPC-21 | IgG1 κ | 551436 | M | BD | matched to test Ab for each lot | | | thresholded in each expt |

*Abbreviations: AF®, Alexa Fluor®; C, citrate buffer; E, EDTA buffer; FACS, fluorescence-activated cell sorting; G, goat; IF, immunofluorescence; IHC, immunohistochemistry; M, mouse; memb, plasma membrane staining; nuc, nuclear staining; WB, Western blot.*

##### *Methylation array profiling & ChIPseq meta-analysis*

DNA was extracted from FACS-sorted hMEC samples, then analysed using Illumina Infinium Omni2.5 arrays in parallel with a familial breast tumour cohort[9]. Histone modification ChIP-seq data were obtained from Pellacani et al[10]. Briefly, bigwig format files were retrieved from www.epigenomes.ca, and mean signal/bin plotted across the region chr22:38365030-38396083 for each histone mark in each cell type.

##### *Analysis of SOX10 expression in cell lines*

We used lines from our in-house bank[11] as well as two primary melanoma cell lines donated by Chris Schmidt (QIMR Berghofer), preselected for low and high *SOX10* RNA levels using in-house expression data. RNA and protein were extracted from cells in the exponential phase of growth using standard Trizol and RIPA buffer methods[12]. *SOX10* mRNA was quantified relative to *RPL13A* as previously described[13]. For Western analysis

(MDA-MB-435, HCC1569, HCC38 cells), protein lysates (30 μg) were resolved by SDS-PAGE then SOX10 and β-actin were detected using standard chemiluminescence (Table-M1).

##### *Stable shRNA knockdown of SOX10 in breast cancer cell lines*

Three pre-validated *SOX10*-targeted shRNA constructs, and a non-targeting negative control (NTNC) construct (pLKO.1), were purchased from Sigma-Aldrich (TRCN0000018984, TRCN0000018987, TRCN0000018988, SHC002). Plasmid DNA was isolated from overnight bacterial cultures, then lentiviral particles were produced by triple transient transfection of HEK-293T (human embryonic kidney) packaging cells with one of the four transfer plasmids (pLKO.1-puro; 2 µg), together with companion plasmids encoding lentiviral packaging and replication elements (2 µg pHR'8.2ΔR + 0.25 µg pCMV-VSV-G; donated by Dr Wei Shi, QIMR Berghofer). Virus-containing supernatants (in target cell media) were then collected over the following two days and filtered (0.45 µm). MDA-MB-435 target cells were seeded at $3.1×10^4/cm^2$ in 6-well plates, then after 24-48 h (at ~50% confluence), cells were infected with filtered viral supernatants, supplemented with 1 mg/mL polybrene (Sigma-Aldrich) for 24 h. Stably transduced cells were then selected with 1 µg/mL puromycin (Sigma-Aldrich) for two weeks to eliminate uninfected cells.

##### *Datasets and processing*

TCGA level-3 normalised RNAseq data (*"rnaseqv2 illuminahiseq rnaseqv2 unc edu Level 3 RSEM genes normalized data.data.txt"*) from the Data Analysis Center Firehose (http://firebrowse.org/) were used for all single-gene analyses (Fig-S2a, Fig-S5e; test group stratification for Fig-7c and Fig-S3; SOX10 heatstrips in Fig-3a, Fig-7d, Fig-S6a, Fig-S6c). Scaled estimate columns of the "*rnaseqv2 illuminahiseq rnaseqv2 unc edu Level 3 RSEM genes data.data.txt*" were used for all other algorithmic analyses.

For methylation datasets, TCGA level 3 Illumina HM450k data were downloaded from the National Cancer Institute Genomics Data Commons (GDC) data portal (https://portal.gdc.cancer.gov/) and processed using the ChAMP package[14]. We applied the champ.filter function to remove problematic probes (those mapping to to X/Y chromosomes, mapping to multiple locations, located near a SNP and non-CG probes). Filtered data were normalized using the champ.norm function, according to the Beta-Mixture Quantile (BMIQ) algorithm; an intra-sample normalisation procedure that corrects the bias of type-2 probe values.

Level-4 GISTIC-2 copy-number data for TCGA cases were downloaded from the Data Analysis Center Firehose (http://firebrowse.org/) and used for correlative analyses with no further processing.

To apply tumour purity cutoffs (TCGA cases), we used a consensus measurement of four different purity estimation methods[15]. For cancers not in this study (ESCA, PCPG, STAD, THYM, UVM), a methylation-based purity estimate was used[16]. Specific thresholds are indicated in the respective figure legends.

With permission from the METABRIC data access committee, normalised Illumina HT 12 expression array data were downloaded from the European Genome-phenome Archive (EGAD00010000210-211). For the ICGC RNAseq dataset, normalised data were downloaded as supplementary data[17] and used with no further processing. Mutational signature data (COSMIC, v2 SigProfiler) were downloaded as raw event counts from ref[17] and HRDetect probability scores for these cases from ref[18].

##### *Differential expression analysis of SOX10-high and -low TNBCs* (Fig-S3)

To characterise the transcriptomic phenotype associated with *SOX10* expression in TNBC, we performed differential expression analysis of *SOX10*-high versus *SOX10*-low (median split) TCGA and METABRIC datasets using limma[19] (differential expression was defined by a corrected *p*-value cutoff of 0.01).

##### *Ontology enrichment analyses*

GO term enrichment analysis was performed using the Generic GO term finder hosted by Princeton University (Lewis-Sigler Institute for Integrative Genomics; https://go.princeton.edu). Geneset enrichment analysis (GSEA) was performed using the Prerank function of GenePattern[20] using 1000 permutations. For Fig-S3, GSEA inputs comprised differentially expressed genes ($q \leq 0.01$) ranked by fold-change in each dataset. The input for all other GSEA experiments was whole transcriptome gene lists ranked by a Spearman correlation coefficient. Biological process genesets (Gene Ontology v7.2; geneset size 15-500) were mined for unsupervised analyses, and neural crest genesets for supervised analyses (Table-S11). Datasets and ranking metrics are indicated in the respective figure legends. Normalised enrichment scores (NES) and corrected *p*-values are reported. GeneGo (Metacore® Clarivate Analytics) and Ingenuity® Pathway Analysis (Ingenuity) were also used to analyse pre-ranked gene lists. REVIGO[21] was used to resolve semantic redundancy and identify major themes amongst the enriched terms.

##### *Weighted gene co-expression network analysis (WGCNA) – module identification & validation*

WGCNA is a powerful network analysis tool that identifies groups of transcripts (modules) that fluctuate in a highly coordinated fashion, implying co-functionality[22,23]. First, it iteratively correlates the expression of every pair of transcripts in a test dataset, producing an adjacency matrix. It then converts this to a topological overlap matrix that reflects net connection weight, accounting for both direct connections and the impacts of shared neighbours. In this study we created 'signed' networks, which reflect the overall topological overlap considering both positive and negative correlations. Dynamic module identification and characterisation (derivation of network metrics, sample eigengene values and module preservation in orthogonal datasets, see below) were performed in the R coding environment, and publication-quality figures were prepared from raw datasets using GraphPad Prism, Clustergrammer or Tableau (see Table-S1 for more details including code availability).

Modules were identified using the TCGA RNAseq (n=919 samples after quality filtering) and validated using METABRIC (n=1278; expression array) (Fig-M1, Fig-M2, Table-M2). A consensus set of 8 modules was determined according to satisfactory concordance between these two orthogonal networks and a third generated from the ICGC dataset (n=342; RNAseq). We further validated the 8 consensus modules using preservation analysis on a third breast cancer expression dataset from. For normal breast samples, WGCNA was performed independently on TCGA normal breast samples (n=97 after quality filtering).

Standard WGCNA outputs include the following (raw data in Tables S5-11):

- <u>Module eigengene (ME)</u>: a theoretical gene that is the most strongly connected to all other genes in the module and hence represents net module expression and connectivity. Mathematically, the first principal component of each module's adjacency matrix.
- <u>Module membership & connectivity:</u> Each gene is ascribed k values describing modular and network connectivity (kTotal, kWithin, kOut). These continuous variables are amenable to integrated analysis of overlapping transcriptional programs, utilizing the granularity in expression datasets rather than levelling it as is done when assigning fixed phenotypes or categories. kME correlation and kME p-values describe how tightly individual genes are linked to all other genes within each module.

To identify hub genes (Fig-M3), additional network connectivity & influence measures were calculated for each node in the SOXE-module topological overlap matrix using igraph toolkit functions in R:

- <u>betweenness centrality</u>: betweenness(graph, v=V(graph), directed=FALSE, weights=NULL, nobigint=TRUE, normalized=FALSE).
- <u>eigencentrality</u>: eigen centrality(graph, directed=FALSE, scale=TRUE, weights=NULL, options=arpack defaults).

Finally, we used community detection algorithms[24,25] to examine the substructure of the SOXE-module (MATLAB 2020a), using the adjacency matrix as input. This revealed a hierarchical, sub-modular organisation, and consistently discriminated two partitions (59% and 41% of nodes each). To identify the module 'control centre' and hub genes as points of structural vulnerability, submodule assignment was cross-referenced against clustered Cosine similarity data (Fig-3b, Clustergrammer[26]) with the same input (Fig-S4).

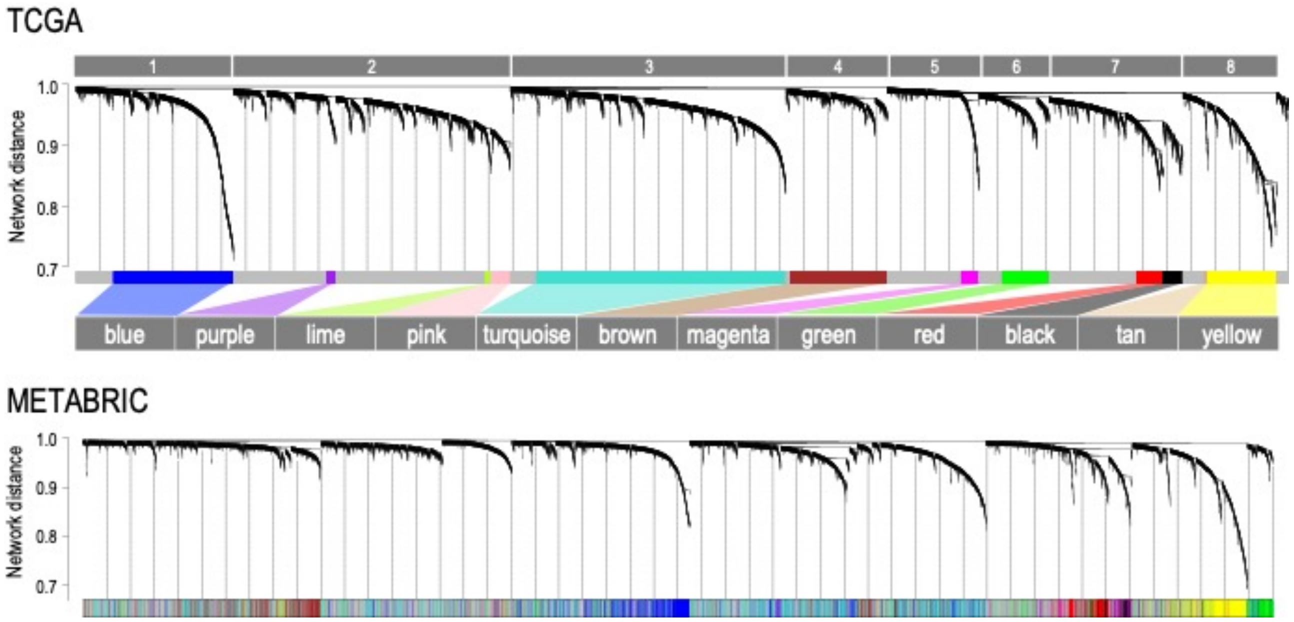


**Figure-M1**: Breast cancer WGCNA network topological overlap visualized after clustering, where lower values on the y-axis indicate progressively shorter inter-gene distance. The TCGA dataset was used for discovery, and METABRIC for validation. After METABRIC module identification, genes were given the same module assignment to highlight overlap in clustering amongst those with the lowest distance.

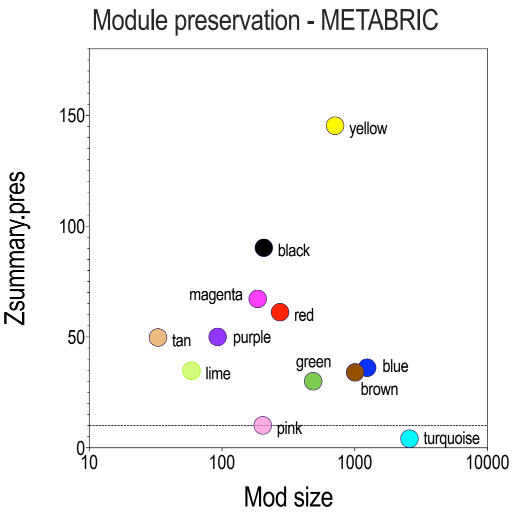


| Module | Module size (no. genes*) | Consensus (kME correlation) | |
|---|---|---|---|
| | | *r* | *p* |
| Blue | 1239 | 0.76 | 1.0E-200 |
| Brown | 1008 | 0.62 | 4.2E-108 |
| Magenta | 186 | 0.80 | 2.6E-39 |
| Green | 487 | 0.61 | 2.6E-43 |
| Red | 274 | 0.50 | 3.9E-13 |
| Black | 207 | 0.56 | 6.7E-14 |
| Tan | 33 | 0.79 | 1.2E-03 |
| Yellow | 712 | 0.70 | 2.7E-83 |
| *Purple* | *93* | *0.33* | *1.7E-04* |
| *Lime* | *59* | *0.47* | *5.1E-03* |
| *Pink* | *203* | *-0.02* | *6.8E-01* |
| *Turquoise* | *2565* | *0.02* | *9.5E-01* |

**Genes correlating with ME with p≤1.0E-05*

**Table-M2** module membership correlation between TCGA and METABRIC datasets. The bottom four modules were filtered out on the basis of low correlation between datasets.

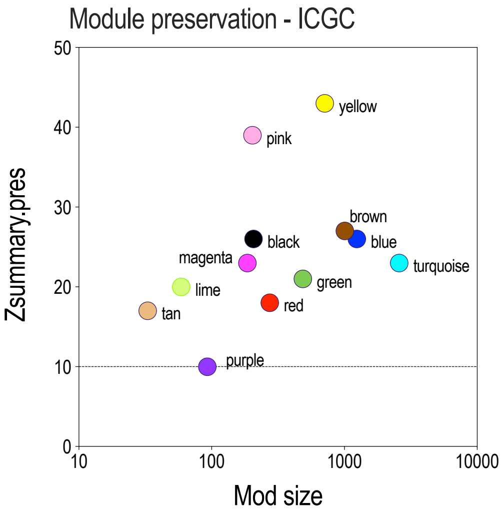


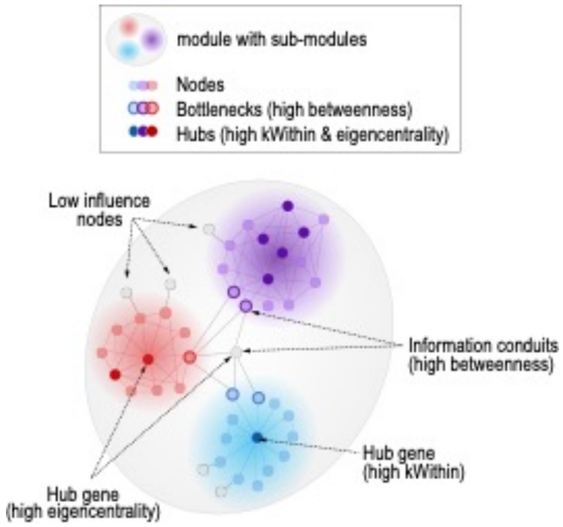


**Figure-M2**: module preservation. Plots: z-score measures of comparison distribution (>10 considered concordant).

**Figure-M3**: Network schematic defining nodes, hubs, modules and node influence metrics.

##### *Neural crest genesets*

Geneset-1 (NC-terms) comprises 308 genes represented in at least two of the 78 terms matching ‘neural crest’ and ‘human’ in the gene ontology database (http://geneontology.org). Geneset-2 (ch.NCSC) comprises the top 200 transcripts statistically over-represented in Sox10+ chick neural crest cells compared to the all other embryo cells (fold-change 3.9–23.3; false discovery rate 9.3E$^{-03}$–1.0E$^{-15}$)[27] (see Table-S11). The ch.NCSC geneset represents genes co-ordinately expressed with *Sox10* in a stem cell state hence was also suitable for network analyses (see below).

##### *Neural crest geneset expression analyses*

We used the *singscore* algorithm[28] to score RNAseq datasets against the neural crest genesets at the individual sample level. To investigate SOX10’s influence over NCSC gene expression, we used *Pathifier*, a principal curve variance-based tool that minimises the impact of small, linear metagenes within larger genesets (in this case, minimising the impact of *SOX10* itself within *SOX10*-stratified test groups)[29-31]. To ensure there was enough variation to support accurate calculation of pathway deregulation scores, the ch.NCSC geneset was extended to include all genes significantly upregulated in Sox10+ chick NCSCs[27] (Table-S11).

##### *Methylation data analyses*

Methylation beta-values were derived from TCGA level 3 Illumina HM450k data as outlined above. Beta-values for all probes corresponding to TSS1500, TSS200 and 5'UTR regions in each sample were first normalized to correct for their bimodal distribution (median absolute deviation (MAD): $P_{\beta}$ – median$(P_{\beta}$ – median$(R_{\beta}))$; where $P$ = probe in the promoter region, and $R$ = all probes in promoter region). After filtering out genes with >2 missing probes and those for which >2% of samples were missing data, the final dataset included average MAD-normalized promoter methylation beta-values for 4482 genes (determined from a total of 518 samples with complete clinical annotation). Pairwise Spearman correlations were then calculated between each promoter region and each module eigengene across the sample cohort. Unsupervised hierarchical clustering of correlation values was performed in R using the *Flashclust* package based on the Euclidean distance method. Clusters were visualised and validated with the *cluster* package, which uses the 'Silhouette Coefficient' to measure how similar a sample is to its cluster compared to a neighbouring cluster. To generate t-distributed stochastic neighbour embedding (t-SNE) plots, we used the Rtsne package (https://cran.r-project.org/web/packages/Rtsne/) on normalized beta methylation values, with 5000 iterations and a perplexity parameter of 40.

##### *TCGA cancer-specific network construction and connectivity analyses*

First, we built Pearson correlation matrices from RNAseq datasets of 22 solid cancer types, then iteratively quantified the pairwise correlation coefficients of sets of 60 genes, selected either at random from NCSC genesets or the entire transcriptome. We then analysed the distributions of median correlation values from 1000 permutations, for three genesets across all cancer types. Random permutation distribution data were used to characterise the transcriptome network structures of each cancer type (Fig-S6b, Fig-S6f), and also as a control with which to compare to neural crest genesets (Fig-S6c, Fig-S6d). Cumulative distributions were compared in GraphPad Prism using the Kolmogorov-Mirnov (K-S) D statistic.

For greater stringency, we then applied a novel approach to strip the correlation matrices of noise. By numerically calculating the equivalent Marchenko-Pastur distribution for the given observational dataset, eigenvalues indiscernible from random noise were able to be removed. This was performed by performing eigendecomposition on the correlation matrices and setting any eigenvalues that lied within the Marchenko-Pastur distribution to zero (0). The correlation matrix was then reconstituted using the cleaned eigenvalues and eigenvectors from the eigendecomposition process. We then derived the following metrics for each malignancy: (1) median eigenvectors of all ch.NCSC gene pairs (intra-geneset connectivity); and (2) median eigenvectors of any gene pair involving ch.NCSC genes, reflecting connectivity to all other genes (geneset-network connectivity).

#### REFERENCES


1. McCart Reed, A.E., et al., *The Brisbane Breast Bank.* Open Journal of Bioresources, 2018. **5**: p. 5.

2. Raghavendra, A., et al., *Expression of MAGE-A and NY-ESO-1 cancer/testis antigens is enriched in triple-negative invasive breast cancers.* Histopathology, 2018. **73**(1): p. 68-80.

3. Abdel-Fatah, T.M., et al., *SPAG5 as a prognostic biomarker and chemotherapy sensitivity predictor in breast cancer: a retrospective, integrated genomic, transcriptomic, and protein analysis.* Lancet Oncology, 2016. **17**(7): p. 1004-1018.

4. Abdel-Fatah, T.M.A., et al., *Association of Sperm-Associated Antigen 5 and Treatment Response in Patients With Estrogen Receptor-Positive Breast Cancer.* JAMA Netw Open, 2020. **3**(7): p. e209486.

5. McCart Reed, A.E., et al., *Phenotypic and molecular dissection of metaplastic breast cancer and the prognostic implications.* J Pathol, 2019. **247**(2): p. 214-227.

6. Saunus, J.M., et al., *Integrated genomic and transcriptomic analysis of human brain metastases identifies alterations of potential clinical significance.* J Pathol, 2015. **237**(3): p. 363-78.

7. Johnston, R.L., et al., *High content screening application for cell-type specific behaviour in heterogeneous primary breast epithelial subpopulations.* Breast Cancer Res, 2016. **18**(1): p. 18.

8. Smart, C.E., et al., *In vitro analysis of breast cancer cell line tumourspheres and primary human breast epithelia mammospheres demonstrates inter- and intrasphere heterogeneity.* PLoS One, 2013. **8**(6): p. e64388.

9. Nones, K., et al., *Whole-genome sequencing reveals clinically relevant insights into the aetiology of familial breast cancers.* Ann Oncol, 2019. **30**(7): p. 1071-1079.

10. Pellacani, D., et al., *Analysis of Normal Human Mammary Epigenomes Reveals Cell-Specific Active Enhancer States and Associated Transcription Factor Networks.* Cell Rep, 2016. **17**(8): p. 2060-2074.

11. Saunus, J.M., et al., *Multidimensional phenotyping of breast cancer cell lines to guide preclinical research.* Breast Cancer Res Treat, 2018.

12. Momeny, M., et al., *Heregulin-HER3-HER2 signaling promotes matrix metalloproteinase-dependent blood-brain-barrier transendothelial migration of human breast cancer cell lines.* Oncotarget, 2015. **6**(6): p. 3932-46.

13. Vargas, A.C., et al., *Gene expression profiling of tumour epithelial and stromal compartments during breast cancer progression.* Breast Cancer Res Treat, 2012. **135**(1): p. 153-65.

14. Tian, Y., et al., *ChAMP: updated methylation analysis pipeline for Illumina BeadChips.* Bioinformatics, 2017. **33**(24): p. 3982-3984.

15. Aran, D., M. Sirota, and A.J. Butte, *Systematic pan-cancer analysis of tumour purity.* Nat Commun, 2015. **6**: p. 8971.

16. Zheng, X., et al., *Estimating and accounting for tumor purity in the analysis of DNA methylation data from cancer studies.* Genome Biol, 2017. **18**(1): p. 17.

17. Nik-Zainal, S., et al., *Landscape of somatic mutations in 560 breast cancer whole-genome sequences.* Nature, 2016.

18. Davies, H., et al., *HRDetect is a predictor of BRCA1 and BRCA2 deficiency based on mutational signatures.* Nat Med, 2017. **23**(4): p. 517-525.

19. Ritchie, M.E., et al., *limma powers differential expression analyses for RNA-sequencing and microarray studies.* Nucleic Acids Res, 2015. **43**(7): p. e47.

20. Subramanian, A., et al., *Gene set enrichment analysis: a knowledge-based approach for interpreting genome-wide expression profiles.* Proc Natl Acad Sci U S A, 2005. **102**(43): p. 15545-50.

21. Supek, F., et al., *REVIGO summarizes and visualizes long lists of gene ontology terms.* PLoS One, 2011. **6**(7): p. e21800.

22. Zhang, B. and S. Horvath, *A general framework for weighted gene co-expression network analysis.* Stat Appl Genet Mol Biol, 2005. **4**: p. Article17.

23. Langfelder, P. and S. Horvath, *WGCNA: an R package for weighted correlation network analysis.* BMC Bioinformatics, 2008. **9**: p. 559.

24. Blondel, V.D., et al., *Fast unfolding of communities in large networks.* Journal of Statistical Mechanics: Theory and Experiment, 2008. **2008**(10): p. P10008.

25. Lambiotte, R., J.C. Delvenne, and M. Barahona, *Laplacian Dynamics and Multiscale Modular Structure in Networks.* arXiv:0812.1770, 2009.

26. Fernandez, N.F., et al., *Clustergrammer, a web-based heatmap visualization and analysis tool for high-dimensional biological data.* Sci Data, 2017. **4**: p. 170151.

27. Simoes-Costa, M., et al., *Transcriptome analysis reveals novel players in the cranial neural crest gene regulatory network.* Genome Res, 2014. **24**(2): p. 281-90.

28. Foroutan, M., et al., *Single sample scoring of molecular phenotypes.* BMC Bioinformatics, 2018. **19**(1): p. 404.

29. Drier, Y., M. Sheffer, and E. Domany, *Pathway-based personalized analysis of cancer.* Proc Natl Acad Sci U S A, 2013. **110**(16): p. 6388-93.

30. Livshits, A., et al., *Pathway-based personalized analysis of breast cancer expression data.* Mol Oncol, 2015. **9**(7): p. 1471-83.

31. Huang, S., et al., *A novel model to combine clinical and pathway-based transcriptomic information for the prognosis prediction of breast cancer.* PLoS Comput Biol, 2014. **10**(9): p. e1003851.
